# Transcriptional Mapping of Neuro-Immune Interactions during Homeostasis and HIV infection using Microglia-containing Human Cerebral Assembloids

**DOI:** 10.64898/2026.08.10.743763

**Authors:** Sheetal Sreeram, Ya Chen, Luke A D Bury, Konstantin S Leskov, Fengchun Ye, Yoelvis Garcia-Mesa, Benjamin Luttge, Jaejin Eum, Junyou Huang, Asha Kallianpur, Anthony Wynshaw-Boris, Jonathan Karn

## Abstract

**Background:** A significant number of people with HIV-1 still experience neurocognitive impairments (NCI), despite effective antiretroviral treatment. HIV-NCI is diverse and multifactorial, with mechanisms that cause its development and progression still not fully understood. We examined early HIV-related changes in brain stability and studied neuroimmune interactions at the single-cell level to better understand how NCI develops.

**Methods:** To model changes in brain homeostasis, we developed an advanced human iPSC-derived 3D cerebral assembloid model that includes microglia, by co-developing neural progenitor cells with tdTomato-tagged and CD34+ cell-derived microglial precursors. Assembloids were infected with a macrophage R5-tropic HIV-1 strain NL-AD8. Viral spread was measured using a proviral DNA assay, qPCR for HIV RNA, and 3D immunostaining for Tat protein. Single-cell transcriptomics with tdTomato lineage tracing revealed HIV-1 induced disturbances and cell-type-specific responses. The niche net algorithm was used to identify ligand-receptor interactions between microglia and the brain microenvironment during homeostasis and HIV infection.

**Results:** Highly ramified tdTomato+ IBA-1+ microglia were evenly distributed throughout the assembloids within 15 days of culture. Single-cell transcriptomics identified microglia, excitatory/inhibitory neurons, astrocytes, and oligodendrocyte precursors within the assembloids. Neurons in microglia-containing assembloids upregulated genes related to neurotransmission, synaptogenesis, and neuronal development compared to neurons in organoids without microglia. Niche net analysis showed microglia-derived neurotropic ligands supported neuronal and astrocytic differentiation. The R5-tropic HIV-1 specifically targeted microglia, inducing a reactive phenotype that transmitted interferon and pro-inflammatory signals to nearby cells and increased MHC-I antigen-presentation genes. Notably, neuroprotective ligands from non-glial cells and bystander microglia in the assembloids attempted to counteract HIV-related inflammation and promote neural repair.

**Conclusions:** Our microglia-containing assembloid model replicates in vivo neurodevelopmental interactions, allowing high-resolution analysis of homeostatic and HIV-induced responses across different brain cell types. Homeostatic microglia support neuronal health, while HIV infection triggers a reactive state that spreads inflammatory signals within the brain environment. The presence of multiple glial and non-glial populations uncovered previously unknown crosstalk, including bystander microglial phenotypes and neuroprotective signaling mechanisms that counteract inflammation. These findings emphasize early HIV responses that balance injury and adaptation, offering insights for developing therapies that target microglial activation, boost neuroprotection, and address HIV reservoirs in the brain.

## Background

At least one-third and as many as half of people with HIV (PWH) experience HIV-associated neurocognitive impairment (NCI) [1, 2], which adversely affects their functional status and quality of life, and reduces adherence to Antiretroviral Therapy (ART) [1, 3–9]. NCI is clinically diverse, and even asymptomatic NCI raises the risk of developing symptomatic cognitive impairment [10].

Although ART significantly reduces HIV RNA levels in the brain [11, 12], ART has not decreased the prevalence of NCI [13, 14]. Demyelination or damage to myelinated (white matter) tracts in the brain occurs early after HIV infection, coincides with the onset of NCI, and continues even after viral suppression, along with subsequent gray matter abnormalities. [15, 16]. Virally suppressed PWH with NCI mainly show synaptodendritic simplification rather than neuronal loss [3, 15, 17]. This indicates that HIV persistence in the central nervous system (CNS) causes subtle but important disruptions in metabolic, neurotrophic, antioxidant, and other supportive intercellular communications, leading to cognitive decline.

Microglia form the initial immune barrier in the brain and serve multiple roles during brain development [18], such as promoting the formation of neuronal synapses through direct cell-to-cell contact and indirectly by secreting growth factors, neural patterning factors, and cytokines [19–27]. In turn, neurons, astrocytes, and non-glial cells actively regulate microglial function and alter CNS inflammatory responses by releasing specific signals, which keep microglia in either a homeostatic state or a reactive phagocytic state [23, 28–35]. The modulation of microglia by astrocytes, neurons, and *vice versa* begins early in development and continues throughout life, maintaining brain homeostasis and limiting microglia-induced neuroinflammation [36].

HIV-1 enters the brain early during peripheral infection (within 5-7 days) and crosses the blood-brain barrier (BBB), either by a Trojan horse mechanism through infected monocytes or lymphocytes, or as free virions [37]. Once the virus enters the brain parenchyma, CNS resident immune cells, such as perivascular macrophages and resident microglia, become infected and serve as long-lived cellular reservoirs of HIV-1, even in virally suppressed individuals [38, 39]. Although astrocytes are the most abundant glial cell type in the human brain and have been shown to harbor HIV-1 DNA *ex vivo* [40, 41], their role in maintaining a persistent CNS reservoir for replication-competent virus remains uncertain [42].

In the CNS microenvironment, neurons and astrocytes support microglia homeostasis, which can help maintain HIV silencing and persistence [43–48]. Imbalances in the homeostatic crosstalk during viral persistence can trigger neuroinflammation, increase oxidative stress, disrupt synapses, and activate microglial cells, all of which contribute to various neurodegenerative diseases and play a crucial role in the development of NCI. [42, 43, 49–53].

Molecular studies of HIV in the brain are limited by the scarcity of available tissue, creating an urgent need for more realistic human brain models for neuroHIV that can replicate in vivo responses to infection and inflammation, such as cerebral organoids (CO) [54]. CO derived from embryonic stem cells (ESCs) or induced pluripotent stem cells (iPSCs) has become an exciting tool because they develop authentic brain-resident cells (neurons, astrocytes, oligodendrocytes, and microglia) within a three-dimensional (3D) structure and mimic many of the complex in vivo interactions between cell types in the CNS. [54–56]. Since CO does not produce microglia, adding microglia to this system to create cerebral assembloids (CA) is essential for studying how HIV-1 infection affects neuro-immune interactions.

Diverse experimental strategies have been employed to integrate microglia into cerebral organoids for modeling CNS HIV infections [49, 56–61]. Boreland et al. generated iPSC-derived microglia in two-dimensional (2D) culture and integrated them into sliced neocortical organoids to examine gene expression changes during HIV infection, mainly using bulk RNA sequencing and qPCR [57]. Kong et al. used choroid plexus organoids containing microglia to investigate the effects of HIV infection; their primary focus was on late-stage neuroinflammation caused by choroid cell populations [60].

In a significant recent study, Narasipura et al. created microglia-containing cerebral organoids to examine global transcriptomic changes after HIV infection. A limitation of their experimental approach was that they could not determine how each cell type was affected by the virus. [61]. In a complementary study, Martinez-Meza et al. used a coculture method where fully differentiated microglial cells were infected with HIV before being integrated with 3D brain organoids. This design allowed the study of neuroinflammatory responses during HIV and ART interruption. However, since the microglia were pre-infected, they were unable to study the impact of early viral entry mechanisms and the complex cell-cell and neuroimmune interactions that initiate HIV pathogenesis. [58]. Thus, despite these experimental advances, significant challenges remain in replicating the homeostatic interactions of the brain microenvironment during both microglia development and HIV infection, as well as in fully capturing the diversity of cell-cell interactions needed to study the onset and severity of HIV-associated NCI.

We adapted an attractive approach described by Xu et al. [62] to generate CA by co-culturing myeloid precursors with neural progenitor cells (NPCs) derived from iPSCs. This method allows the differentiation of microglia within the assembloids alongside neural progenitor-derived non-glial cells, recapitulating normal neurogenesis. Using these 3D cerebral assembloids containing microglia and single-cell RNA-sequencing (scRNA-seq) analysis, we identified perturbations caused by HIV infection in the assembloids. Consistent with previous reports [49, 57, 58, 60, 61, 63], we found activation of interferon and proinflammatory signaling in HIV-infected (HIV+) microglia very early during infection. However, because multiple glial cells are resident in the assembloids, we could also define homeostatic crosstalk between brain cell types, resulting in a better characterization of the complex interplay between neuroinflammatory and neuroprotective signaling in the assembloid microenvironment.

## Results

### Co-development of Microglia precursors and Neural progenitors in 3D human cerebral assembloids

**Fig. 1** illustrates our method for generating cerebral assembloids containing microglial cells (MG-CA). We mimicked the early events of neurogenesis by allowing primitive myeloid precursors to develop alongside neural progenitors, as described by Xu et al. [62]. To track microglia migration and differentiation in the assembloids, we produced microglia precursors using the control COR-LT (Clay) iPSC line with a permanently activated nuclear-localized tdTomato (tdT) expression, as described in Bury et al. [64]. To form MG-CA, tdT-negative NPCs were combined with tdT-tagged CD34+ CD45+ hematopoietic (myeloid) precursors on day 1 in a 7:3 ratio and cultured as 3D embryoid bodies with microglia growth factors IL-34 (100 ng/mL), TGFβ1 (50 ng/mL), and M-CSF (25 ng/mL) (**Fig. 1A**). The presence of tdT allowed visualization of CD34+ cells in assembloids throughout their differentiation into microglia. We also created organoids with only NPCs (10:0 NPC:CD34+ ratio) as control CO without microglia (**Fig. 1A**). On day 15, both MG-CAs and CO were transferred to neural maturation media containing growth factors like brain-derived neurotrophic factor (BDNF), glial cell line-derived neurotrophic factor (GDNF), cyclic-AMP (cAMP), and ascorbic acid to promote neural maturation, along with IL-34, TGFβ1, and M-CSF for microglia differentiation and survival. We monitored the organoids and assembloids in culture for 25-30 days and conducted all experiments within this period.

**Figure 1.**
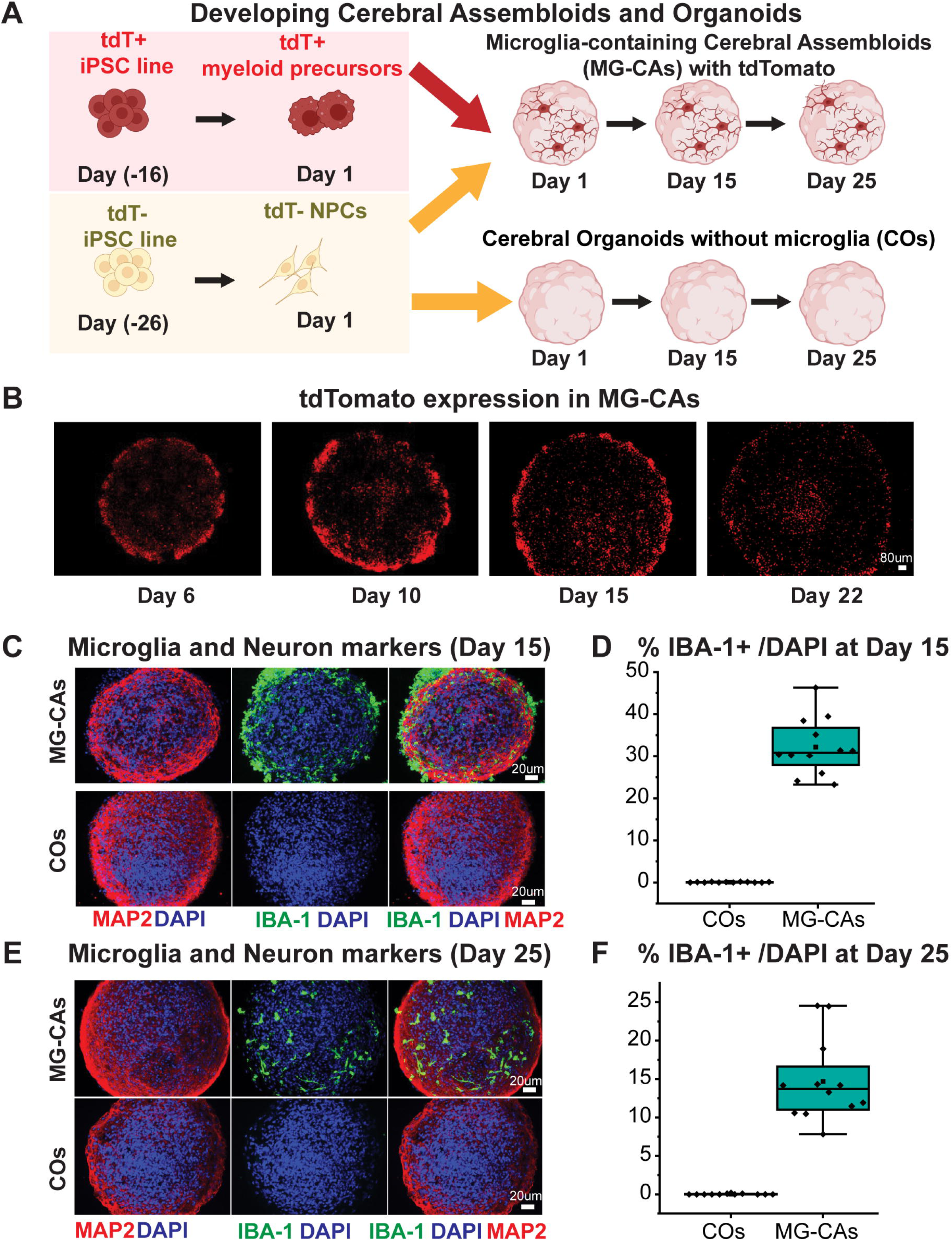
Uniform distribution of IBA1+ microglia in assembloids. **A)** Schematic for deriving microglia-containing cerebral assembloids (MG-CAs) by co-culture of NPCs and tdTomato+ microglial precursors and cerebral organoids (CO) lacking microglia from tdTomato-NPCs. Image created using BioRender. **B)** Live imaging of tdTomato protein expression in MG-CAs from weeks 1-3 (day 6 to day 22) post assembloid formation. **C)** Representative images showing IBA-1 (Microglia marker, green) and MAP2 (Neuronal marker, red) staining in MG-CAs and COs at Day 15 of cultures. **D)** Bar graphs depicting percent IBA-1+ cells in COs and MG-CAs at Day 15 post culture. N=12 organoids or assembloids, p < 0.001. **E)** Representative images showing IBA-1 (Microglia marker, green) and MAP2 (Neuronal marker, red) staining in MG-CAs and COs without Microglia at Day 25 of cultures. **F)** Bar graphs depicting percent IBA-1+ cells in COs and MG-CAs at Day 25 post culture. N=12 organoids or assembloids, p < 0.001. Scale bars, as described in images.

The co-development of tdT-tagged CD34+ CD45+ myeloid precursors with NPCs enabled improved tdT+ cell migration and homing into the brain assembloids. Microglia integrated throughout the assembloids could be observed as early as one week after coculture (**Fig. 1B**). At earlier time points, we mainly saw tdT+ cells located along the edges of the assembloids. While some tdT+ cells at the edges might have been lost over time due to media changes, the remaining cells gradually migrated inward.

Significant modifications we made included partially differentiating the NPC and microglia ex vivo before assembling the assembloid. Specifically, to generate CD34+ hematopoietic stem cells (iHSC) from iPSCs, we used a serum-and feeder-free 14-day differentiation protocol from iPSCs with the STEMDiff iHSC kit (STEMCell Tech), which mimics the early primitive hematopoietic precursors’ development in the yolk sac (YS) [65, 66]. The iHSC population on day (-4) of differentiation showed strong tdT expression, with over 90% of cells being CD43+ CD34+ double-positive, a marker of hematopoietic progenitor-like cells (**Supplementary Table 1**). We further directed the iHSCs to become microglial precursors by culturing the day (-4) iHSCs for 5 days in microglia medium containing neuron-and astrocyte-secreted factors, IL-34, TGFβ1, and M-CSF. This process mimics primitive microglia differentiation during their migration toward the neural tube [67, 68] and led to a gradual decrease in hematopoietic CD34+ marker (Supplementary Table 1), due to myeloid lineage commitment [69]. NPCs were differentiated from the untransfected control COR-LT (Clay) iPSCs line (minus tdT) using PSC Neural Induction Medium (Gibco) from iPSCs for 7 days and then expanded up to passage 4 for 28 days, as described in the methods. The identity of the NPCs was confirmed by staining for NESTIN, a marker for NPCs, as well as SOX2 and PAX6, before organoid culture.

To monitor microglia and neural markers simultaneously using immunofluorescence, we also used a second healthy control iPSC line lacking tdT (CS00iCTR-n2, obtained from the Cedars-Sinai iPSC core) to derive CD34+ CD45+ hematopoietic (myeloid) precursors and NPCs. At day 15 of coculture, we observed IBA-1+ microglial cells developing in our assembloids, which accounted for approximately 30% of the total DAPI+ cells and MAP2+ young neuronal populations, indicating differentiation of both microglial and neuronal cell types **(Fig. 1C-D, Supplementary Table 1)**. By day 25, IBA-1+ cells had become more dispersed within the assembloids. Due to the reduced number of cells on the surface of the assembloid, the total IBA-1+ levels dropped to about 10-15% of total cells by day 25 **(Fig. 1E-F, Supplementary Table 1)**. Nevertheless, the level of microglia in our model stayed within the 5–20% range, which matches the percentage of microglial cells in the human brain [70–72]. In contrast, no IBA-1+ cells were observed in CO formed from NPCs alone, indicating that spontaneous microglial differentiation did not occur in the organoids.

In summary, by expanding both CD34+ microglial precursors and NPC cultures in their respective media before generating organoids and assembloids, we were able to accelerate the differentiation of both glial and non-glial populations and produce MG-CAs and COs with significant neuronal differentiation even by day 15 of culture.

### Transcriptomic characterization of assembloid microglia

To rigorously define the diversity of cell types in our MG-CAs and COs, as well as the status of microglial differentiation, we performed single-cell RNA sequencing on day 15 of co-culture by pooling approximately 20 organoids and assembloids from both tdT+ MG-CAs and COs. We identified 10 distinct molecular clusters through unsupervised clustering using the Louvain method, after filtering out low-quality cells [73, 74] **(Fig. S1A)**. Clusters 3, 5, 6, 7, 9, and 10 were exclusively found in MG-CAs compared to COs. Clusters 3, 9, and 10 were identified as microglia based on mapping to the Spatio-Temporal cell Atlas of the human Brain (STAB) [75], while Clusters 5, 6, and 7 mapped to NPCs and other neuronal subtypes **(Fig. 2A)**. The tdT RNA+ cells primarily formed a distinct cluster on the UMAP. They were highly enriched in the microglia cluster, confirming the myeloid origin of these cells **(Fig. 2B)**.

**Figure 2:**
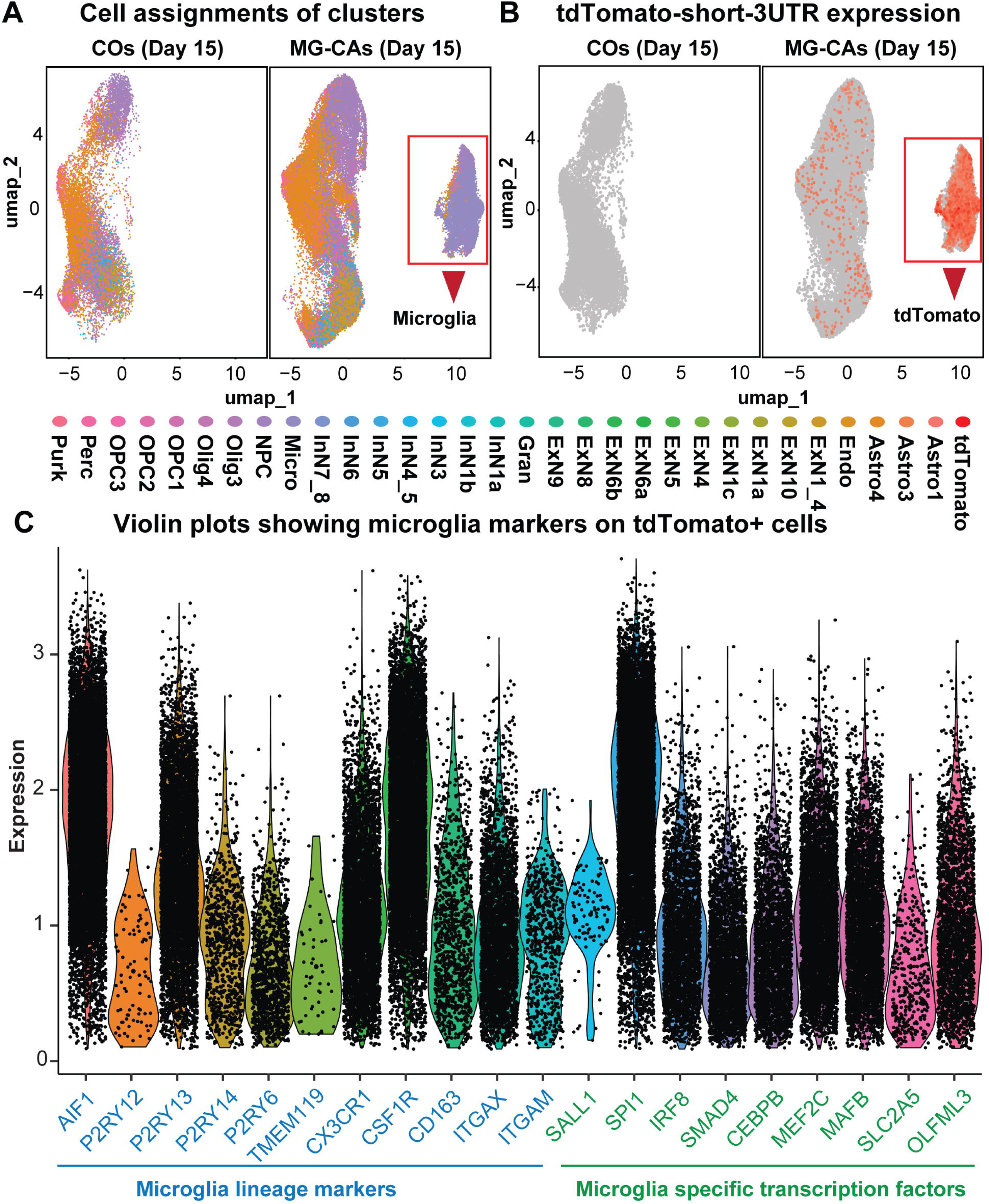
Transcriptomic characterization of tdTomato+ cells in assembloids reveals distinct microglial signatures. **A)** UMAP plots showing the mapping of assembloid and organoid cells from Day 15 of culture onto the Spatio-Temporal Atlas of the Human Brain (STAB) reference dataset. Plots are split by sample type: COs and MG-CAs. **B)** Feature plots displaying tdTomato RNA expression across MG-CAs and COs. **C)** Feature plots showing expression of key homeostatic microglia markers; P2RY13 (purinergic receptor), CX3CR1 (fractalkine receptor), AIF1 (also known as IBA1), and CSF1R (colony-stimulating factor 1 receptor) in the tdTomato+ cluster from **(B)**.

To further examine microglial differentiation and phenotypic states in the assembloids, we subgrouped the tdT+ cells (tdT RNA reads > 0) from **Fig. 2B** and performed an unsupervised clustering analysis. We identified 7 transcriptionally distinct subclusters within the microglial tdT+ population using the Louvain method (**Fig. S1B**). All tdT+ subclusters expressed the homeostatic microglia markers P2RY12, P2RY13, CX3CR1, AIF1, TMEM119 and CSF1R (**Fig. 2C**). However, there were decreased levels of some key homeostatic and mature markers, such as P2RY12 and TMEM119, in the assembloid microglia (**Fig. 2C**). The reduced expression of mature microglial markers in 3D assembloids may be due to the absence of vasculature, oxygenation, and in vivo microenvironment, as noted by Schafer et al. [76] and others [18, 30, 77, 78]. However, microglia from assembloids are capable of further differentiation, and mature markers can be induced by transplantation of assembloids into a mouse brain [76, 79–81]

Nevertheless, by profiling the top markers in each of the tdT+ sub-clusters from **Fig. S1B (Fig. S1C, Supplementary Table 2)**, we observed proliferative microglia (TOP2A, MKI67 in subcluster 3) with self-renewal capacity, which has been previously observed in human primary microglia [82]. Interestingly, tdT+ sub-clusters 6 and 7 expressed complement, phagocytosis genes (CD163, C1QB, C5AR1), chemo-sensing, and cytokine response genes (e.g., P2RY8, CCR7, IL7R), which are necessary for microglial synapse pruning and immune functions [82–84]. tdT+ sub-clusters 0, 1, 2, 4, and 5 exhibited neurodevelopmental markers such as NSG2, NAV3, NRG3, and POU3F1, indicating that assembloid microglia can express neurodevelopmental genes [85–87]. The ability of microglia to express genes involved in neurogenesis and neurodevelopment has been previously observed by Grassivaro et al. [88], who demonstrated that these markers appear during early embryonic development (E14.5) and early postnatal stages (P1) in mouse microglia. Therefore, we observed a highly diverse microglia population in the assembloid at these early stages of differentiation and development, which aligns with previous reports by Popova et al. [82] and others [18, 30, 81, 82, 89].

### HIV productively infects and spreads within microglial populations in cerebral assembloids

To evaluate the impact of HIV-1 infection on cerebral organoids and assembloids, we infected both CO and MG-CAs with the replication-competent, macrophage R5-tropic NLAD8 HIV-1 at Day 15 of culture, after transferring the organoids and assembloids to neural maturation media **(Fig. 3A)**. To assess viral spread, we analyzed the samples from 3 to 6 days post-infection (dpi) using the potentially intact proviral load assay (PIPL) [90] and RT-dPCR for HIV mRNA [91].

**Figure 3:**
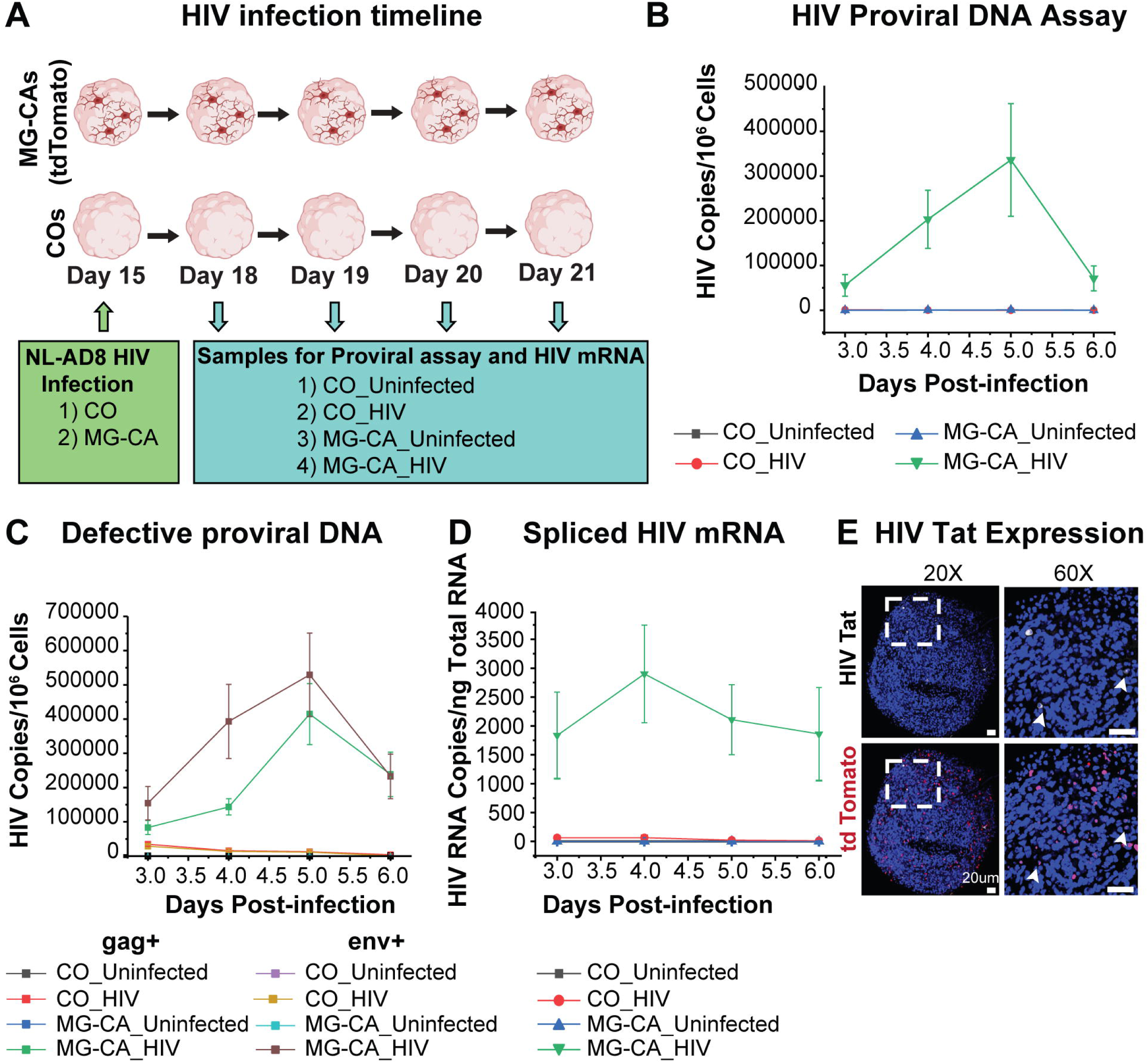
Microglia containing assembloids harbor replicative HIV. **A)** Experimental timeline for HIV infection of organoids and assembloids with the macrophage-tropic NLAD8 replicative HIV strain, followed by detection of HIV proviral DNA and spliced mRNA using digital PCR. Quantification of viral spread in COs and MG-CAs from 3 days post-infection (dpi) to 6 dpi. Image created using BioRender. Plots **B)** show levels of intact proviral DNA (gag+ env+) and **C)** show levels of defective proviral DNA (gag+ or env+) from 3dpi to 6dpi, with n = 5 replicates from two independent iPSC donor lines. Error bars represent Mean ± SEM **D)** Quantification of spliced HIV transcript levels in COs and MG-CAs from 3 dpi to 6 dpi with n = 5 replicates from two independent iPSC donor lines. Error bars represent Mean ± SEM **E)** 3D immunostaining of HIV-infected MG-CAs showing HIV-Tat protein expression and colocalization with tdTomato reporter expression.

The PIPL assay is a two-color digital PCR measurement that uses primers and probes targeting the gag and env regions of the viral genome. This enables measurement of the proportion of double-positive proviral DNA, which is enriched in intact genomes, and single-positive proviral DNA, which may have deletions that can occur during viral reverse transcription. As expected for a productive infection, HIV-1 NL-AD8 infected MG-CAs (MG-CA_HIV) showed increased levels of intact gag+ env+ proviral DNA by 3 dpi. In contrast, no detectable intact proviruses were seen in HIV-infected CO (CO_HIV) **(Fig. 3B-C, Supplementary Table 3)**. In the complementary RT-dPCR assays for HIV-1 mRNAs, we observed an increase in copies of spliced HIV mRNA products from 3 dpi to 4 dpi in the MG-CAs. The spliced mRNAs gradually declined by 5 dpi as some cells died and others likely entered latency. Conversely, control organoids lacking microglia showed no detectable proviruses or HIV RNA **(Fig. 3D, Supplementary Table 3)**. To examine HIV-infected cells histologically, we performed 3D immunostaining on MG-CAs_HIV. We found that HIV-Tat protein localized with tdTomato+ cells **(Fig. 3E)**. Therefore, HIV infects and persists exclusively within microglial cells, which are the main cell types acting as HIV reservoirs in our assembloids.

### Enhanced neurodevelopmental signatures in MG-CAs

An unexpected finding was the higher proportion of neurons in MG-CA compared to CO **(Fig. 2A)**. To examine cell-type-specific gene expression more closely, we performed scRNA-seq on Day 18, three days after switching COs and MG-CAs to neural maturation media. We also included scRNA-seq data from both uninfected and HIV-infected COs and MG-CAs, with about 40 organoids per condition across two separate experimental batches **(Fig. S2A)**. Samples from both replicate batches 1 and 2 were combined for downstream analyses. Using the Seurat integrative analysis pipeline, we identified 13 genetically distinct clusters through Louvain clustering. [73, 74] **(Fig. S2A)**. Specific cell identities for each cluster were determined by mapping reads to the STAB human brain reference dataset, as described previously **(Fig. S2B)**, and by analyzing the top differentially expressed genes (DEGs) between each cluster **(Supplementary Table 4)**.

Based on these analyses, we assigned distinct cell types to each of the 13 cell clusters obtained **(Fig. S3C, Fig. 4A)**. Microglial cells, marked by tdTomato, AIF1, and P2RY13 expression, were identified, along with proliferative NPC1 characterized by high expression of MKI67 and TOP2A, and PAX6-marked neurogenic NPC2. Excitatory neurons (Ex-Neurons) expressed synaptic genes SYN1 and NRXN2, whereas inhibitory neurons (Inh-Neurons) expressed GAD1 and GAD2. OLIG1 and OLIG2 expression identified Oligodendrocyte progenitor-like (OPC-like) cells. In contrast, astrocytic cells showed relatively higher expression of GFAP and AQP4 compared to other clusters **(Fig. S2D)**.

**Figure 4:**
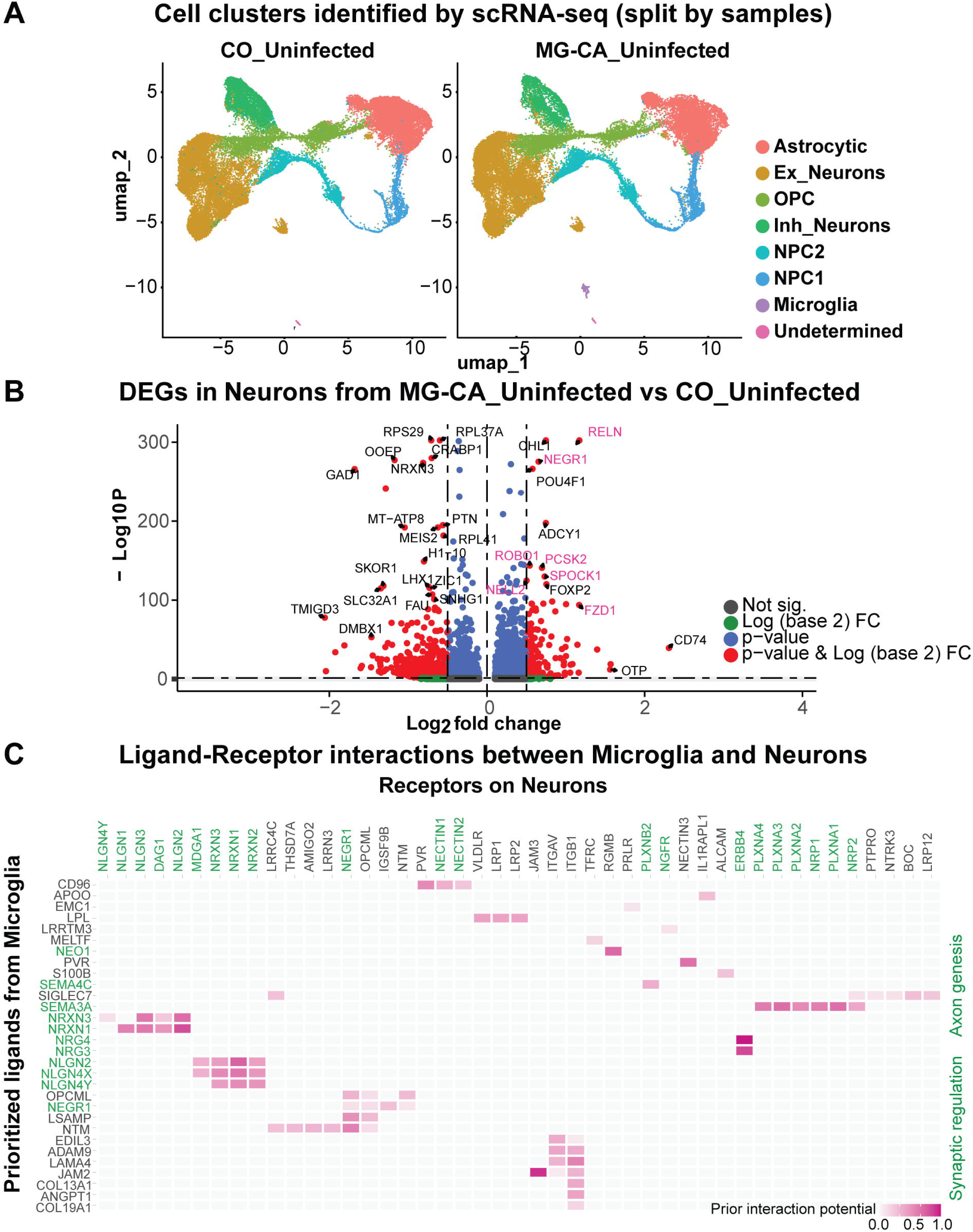
Transcriptomic profiling reveals enhanced neurodevelopmental signatures in MG-CAs compared to COs. **A)** UMAP plots showing unsupervised clustering of cells from CO_Uninfected and MG-CA_Uninfected. Cell type identities were assigned as described in Fig S3 **B)** Volcano plot comparing DEGs between neurons from MG-CAs_Uninfected and CO_Uninfected. The x-axis represents average log2 (fold change), while the y-axis shows-log10 (p-value). Genes with a log2 (fold change) of 0.5 and p < 0.01 are highlighted with red dots. Genes involved in GSEA pathway analysis from **Supplementary Table 4** are highlighted in pink. **C)** NicheNet interactome analysis predicting key ligand-receptor interactions between microglia and neurons (Ex_Neurons and Inh_Neurons). The y-axis represents potential top ligands expressed by microglia, while the x-axis shows corresponding neuronal receptor genes. Interaction strength is represented by color intensity. Ligands involved in synaptogenesis and axon remodeling are labeled in green.

To investigate the role of microglia in assembloid development without HIV infection, we first compared neuronal gene signatures (excitatory and inhibitory) in MG-CA_Uninfected and CO_Uninfected. Gene Set Enrichment Analysis (GSEA) was used to measure DEGs based on log2 fold change. Neurons from MG-CAs showed significant enrichment in pathways related to synaptic assembly, organization, projection, differentiation, and signaling **(Supplementary Table 4)** compared to CO neurons. Using a volcano plot, we identified the top DEGs in neurons from MG-CAs and COs. These included notable upregulation of genes involved in neurotransmission and synaptogenesis (RELN, NEGR1, ROBO1, FOXP2), neuronal development (POU4F1, RELN, NELL2, Wnt pathway receptors like FZD1), and cell-adhesion molecules (CHL1, SPOCK1) in MG-CAs **(Fig. 4B)**. Conversely, genes indicative of early lineage development, progenitor-like states, and reduced synaptic pruning activity, such as GAD1, LHX1, FAU, MEIS2, DMBX1, and OOEP—were among the genes more highly expressed in CO neurons. These findings suggest that microglia accelerate neuronal maturation by promoting synaptic signaling, synapse formation, and neurogenesis in the assembloids.

### Inter-cellular signaling during assembloid development

To better understand cell-cell interactions and study ligand-receptor interactions within the MG-CA microenvironment that promote neuronal maturation and development, we applied the NicheNet algorithm, as described by Saeys et al. [92]. This computational method identifies and ranks ligand-receptor pairs based on the transcriptomic profiles of the ‘sender’ and ‘receiver’ populations, predicting the potential of ligands to induce a set of target genes in ‘receiver’ cells. To compare the microenvironments of MG-CAs and COs in terms of neurogenic conditions, we used the sender-focused approach in NicheNet analysis to predict potential ligand interactions between microglia and neurons based on the expression levels of corresponding receptors and ligands in both sender and receiver cells. CO_Uninfected, which lacks microglia, served as a reference for this analysis.

Our analysis identified several ligand-receptor pairs with high interaction potential involved in microglia-neuron communication. Notable interactions included synaptic guidance ligands, such as Neurexin (NRXN-DAG1, NLGNs), Neuregulin (NRGs-ERBB4), Plexin (PLXN), and cell-adhesion molecules (JAM3, NECTINs), all of which are essential for axon growth, neuronal connectivity, and cell adhesion. Additionally, we observed interactions with neuron growth factor receptors (NGFR1) and other ligands, such as NEO1, which are involved in shaping neuronal synapses **(Fig. 4C)**. These findings highlight the ability of microglia to promote a supportive microenvironment in assembloids that enhances neurogenesis, axonogenesis, glial differentiation, synaptic function, and the maturation of neural networks.

### Enhanced interferon responses in HIV RNA+ Microglia

To characterize gene expression changes in assembloid cell types during early stages of HIV infection, we performed scRNA-seq on MG-CAs at three days post-infection with the NLAD8 strain of HIV. HIV-infected cells within the assembloids and organoids were identified by aligning sequencing reads to the full-length HIV genome [93] using the 7 Bridges platform (BD Biosciences) before Seurat object generation. Increased HIV transcript levels were observed exclusively in the microglial cluster, indicating that microglia are the main cell type harboring HIV in assembloids, aligning with our digital PCR results **(Fig. 5A)**. Analysis of the top DEGs in microglial clusters between MG-CA_HIV and MG-CA_Uninfected showed upregulation of genes related to interferon responses, including MX1, XAF1, IFITM3, OAS3, and IRF9 in microglia from MG-CA_HIV **(Fig. 5B)**. GSEA pathway analysis comparing microglia from MG-CA_HIV and MG-CA_Uninfected further confirmed activation of Type I and II interferon pathways and TNFα-related signaling in microglia from MG-CA_HIV versus MG-CA_Uninfected **(Supplementary Table 5)**.

**Figure 5:**
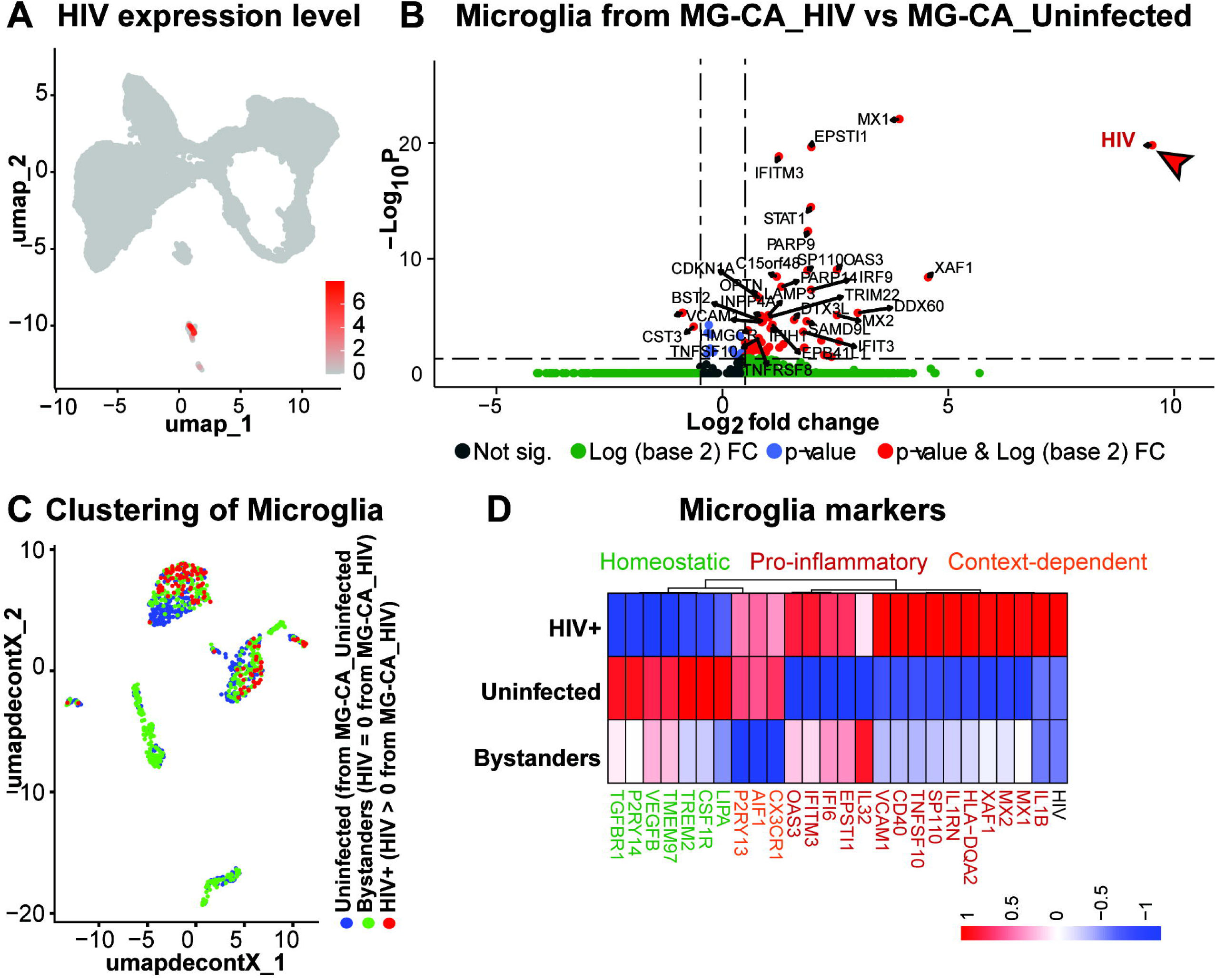
HIV RNA+ Microglia show reactive phenotype compared to bystander and uninfected microglia. **A)** Feature plots showing expression of HIV RNA in the microglial cluster **B)** Volcano plot comparing DEGs between microglia from MG-CAs_HIV and MG-CAs_Uninfected. The x-axis represents average log2 (fold change), while the y-axis shows-Log10 (p-value). Genes with a log2 (fold change) > 1 and p < 0.01 are highlighted with red dots. **C)** UMAP plots showing supervised clustering of microglial cells based on the HIV RNA expression level and categorized as HIV RNA+ microglia (HIV RNA reads > 0 in MG-CA_HIV), bystander microglia (HIV RNA reads = 0 in MG-CA_HIV), and uninfected microglia from MG-CA_Uninfected. **D)** Heatmap showing the average expression of key inflammatory and homeostatic marker genes in the three populations of microglia. Scale shows z-scored average expression, with red indicating high expression, blue indicating low expression. Genes highlighted in red are pro-inflammatory and reactive microglial markers, in green are markers involved in homeostatic responses, and in orange are microglia markers involved in both reactive and homeostatic functions and implicated as per the context.

To analyze the effect of HIV on microglial subpopulations, we divided microglia into three groups based on HIV RNA levels: HIV RNA+ microglia (HIV RNA reads > 0 in MG-CAs_HIV), bystander microglia (HIV RNA reads = 0 in MG-CAs_HIV), and uninfected microglia from MG-CAs_Uninfected **(Fig. 5C)**. Quantifying the phenotypic differences among these microglial populations showed that HIV RNA+ microglia had higher expression of pro-inflammatory genes (IL1B, TNF, IL32, TNFSF10, IL1RN), interferon-stimulated genes (MX1, MX2, OAS3, IFITM3, IFI6, ISG15), and reactive microglial markers (HLA-DQA2, SP110) compared to bystander and uninfected microglia **(Fig. 5D)**. Conversely, homeostatic microglial genes (P2RY13, CSF1R, TGFBR1, CX3CR1, VEGFB) were downregulated in HIV RNA+ microglia.

Bystander microglia showed a partly activated state, with lower levels of inflammatory and interferon gene expression compared to HIV RNA+ microglia, and higher levels of specific homeostatic markers, including VEGFA, TGFBR1, P2RY14, and TREM2 **(Fig. 5D)**. GSEA pathway analysis revealed clear phenotypic differences between bystander and HIV RNA+ microglia. Bystander microglia were enriched for homeostatic pathways, such as synaptic regulation and axon development, while HIV RNA+ microglia were enriched for pathways involved in immune activation and complement signaling **(Supplementary Table 5)**. This analysis highlighted the phenotypic differences between the two microglial cell populations within the MG-CAs_HIV and showed that HIV RNA+ microglia tend to shift toward a highly reactive inflammatory state.

Mapping the autocrine signaling and interactions within microglial clusters showed that HIV RNA+ microglia mainly expressed pro-inflammatory ligands, including TNF, IL1B, HLA-A, and A2M **(Fig. 6A-B)**. These ligands led to the activation of interferon-stimulated genes (IFITs, IRFs) and reactive microglial markers (VCAM1), compared to bystanders and uninfected microglia **(Fig. 6C-D)**. Conversely, bystander microglia expressed higher levels of neuroprotective and immunoregulatory ligands SERPING1 and SIRPA **(Fig. 6B)**. These results suggest that during early HIV replication, HIV RNA+ microglia develop a highly reactive inflammatory phenotype, driving pro-inflammatory signaling and influencing the microenvironment of the assembloid. In contrast, bystander microglia seem less reactive, maintaining homeostatic and neuroprotective roles.

**Figure 6:**
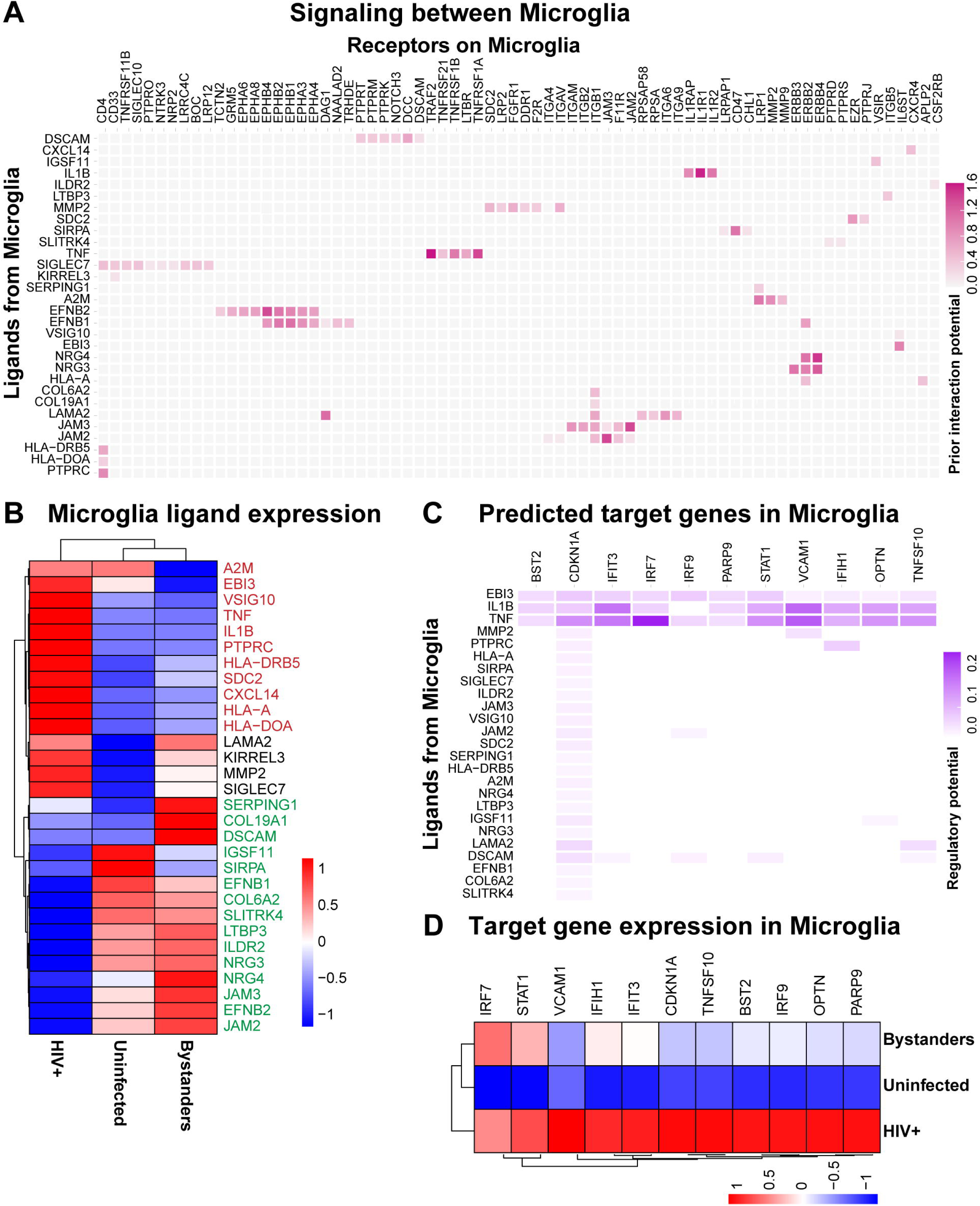
HIV RNA+ Microglia drive reactive states by autocrine interferon and pro-inflammatory ligands. **A)** NicheNet interactome analysis predicting key ligand-receptor autocrine interactions within microglial cells. The y-axis represents potential top ligands expressed by microglia, while the x-axis shows corresponding interactions with receptor genes in microglia. Interaction strength is scaled in pink. **B)** Heatmap shows the average expression of the potential ligands (from panel A) in HIV RNA+, bystander, and uninfected microglia populations. Scale shows z-scored average expression, with red indicating high expression, blue indicating low expression. Genes highlighted in red are pro-inflammatory and reactive microglial ligands, in green are ligands involved in immune-regulatory and homeostatic responses. **C)** NicheNet interactome analysis predicting the regulatory potential of downstream target genes activated in microglia from autocrine signaling. Potential top ligands from microglia are shown on the y-axis and downstream target genes in microglia on the x-axis, with the strength of regulatory potential colored in violet. **D)** Heatmap shows the average expression of the downstream target genes in HIV RNA+, bystander, and uninfected microglia populations. Scale shows z-scored average expression, with red indicating high expression, blue indicating low expression.

To assess how microglial subpopulations affect the MG-CA microenvironment during HIV infection, we mapped potential ligands from microglia to all other cells in the assembloid. In MG-CAs_HIV, pro-inflammatory ligands (TNFSF10, MMP2, HLA-A, CXCL14), along with neuroprotective ligands such as FGF2 [94], ephrin family genes (EFNBP), neuronal guidance ligands (NLGN2, NEGR1, SEMA6C, CNTN2, JAM3, DSCAM), showed higher interaction potential with corresponding receptors on other assembloid cells **(Fig. 7A)**. Comparative analysis indicated that HIV RNA+ microglia mainly expressed pro-inflammatory ligands. Meanwhile, bystander microglia continued to express neuroprotective ligands, maintaining a relatively homeostatic state **(Fig. 7B)**.

**Figure 7:**
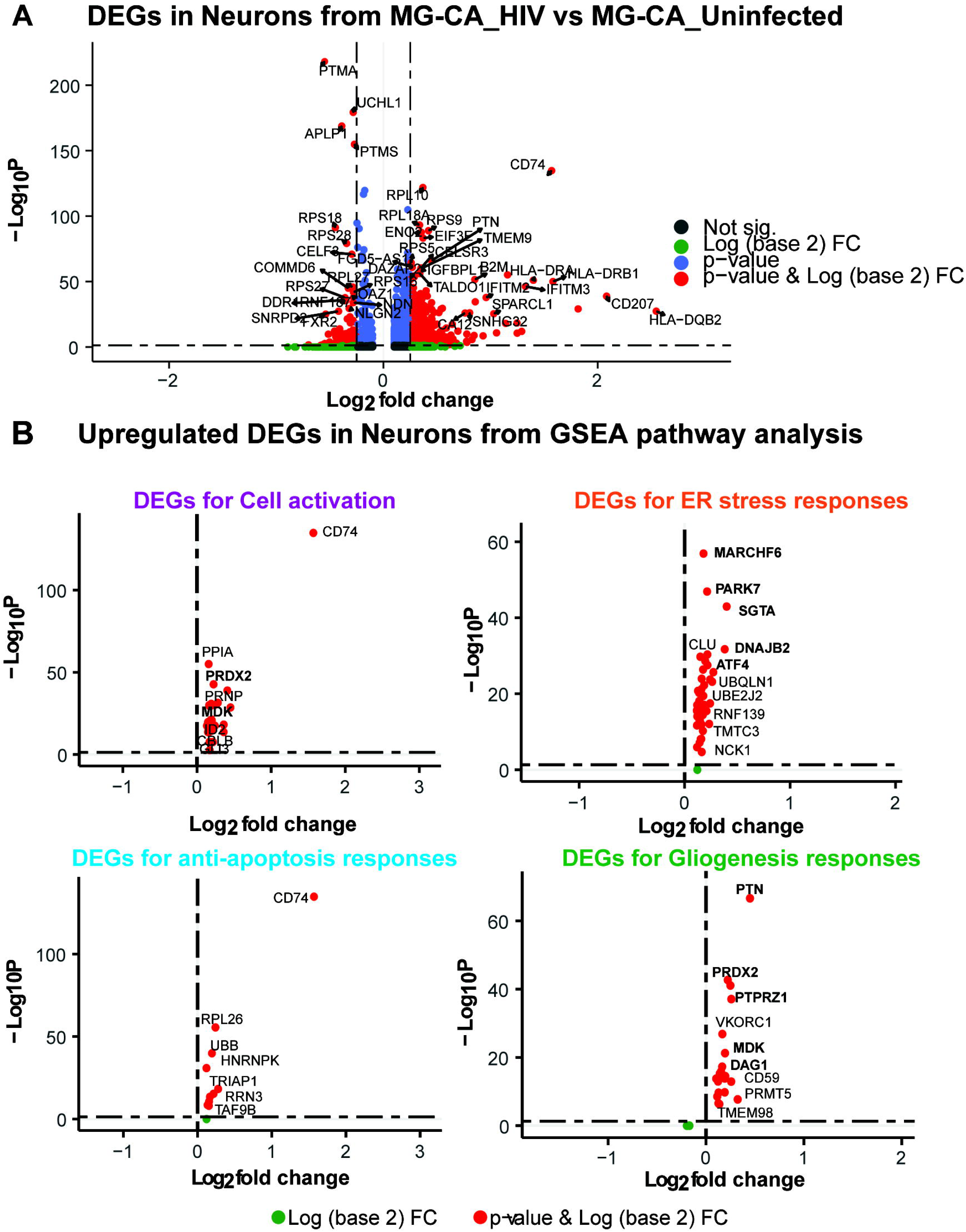
HIV RNA+ Microglia relay pro-inflammatory signals to the assembloid microenvironment compared to bystander microglia. **A)** NicheNet interactome analysis predicting key ligand-receptor interactions between microglia and all assembloid cell types. The y-axis represents potential top ligands expressed by microglia, while the x-axis shows corresponding receptor genes in all assembloid cell types. Interaction strength is represented by color intensity. Ligands involved in pro-inflammatory responses are highlighted in red. **B)** Heatmap shows the average expression of microglia ligands (from panel A) in the HIV RNA+, bystander, and uninfected microglial populations. Ligands expressed in HIV RNA+ microglia are highlighted in red. Scale shows z-scored average expression, with red indicating high expression, blue indicating low expression.

### Pro-inflammatory conditions in HIV-infected assembloids induce stress, repair, and immune-modulatory responses in neurons

Neuronal, astroglial, and other non-glial dysfunctions are known to accelerate the progression of HIV-associated neurocognitive disorders (HAND). We therefore investigated the early cellular effects of HIV at 3 dpi on neurons within the assembloids.

To understand how HIV affects different neuronal subpopulations, we conducted differential expression analysis comparing neurons, both excitatory and inhibitory, from MG-CA_HIV with those from MG-CA_Uninfected. Neurons from MG-CA_HIV showed upregulation of genes related to antigen presentation, such as CD74, B2M, IFITM3, and CD207. Notably, both MHC class I and II genes were elevated, indicating that pro-inflammatory cytokines and interferons during infection likely promoted neuronal peptide processing and antigen presentation. Additionally, we observed increased expression of ribosomal genes (RPL37A, RPL38, RPS21) and genes linked to neural development, including PTN (pleiotrophin), SPARCL1, and IGFBPL1, in neurons from MG-CA_HIV **(Fig. 8A)**. A GSEA pathway analysis was performed to identify pathways disrupted by HIV in neuronal cells. Due to gene overlap across pathways, we combined related pathways sharing similar gene sets **(Supplementary Table 6)**. This grouped analysis highlighted enrichment in four key pathways during HIV infection: 1) cell activation and antigen presentation responses, 2) endoplasmic reticulum (ER) stress responses, 3) gliogenesis and OPC differentiation, and 4) anti-apoptotic responses, suggesting disruption of neuronal homeostasis during HIV infection **(Supplementary Table 6)**.

**Figure 8:**
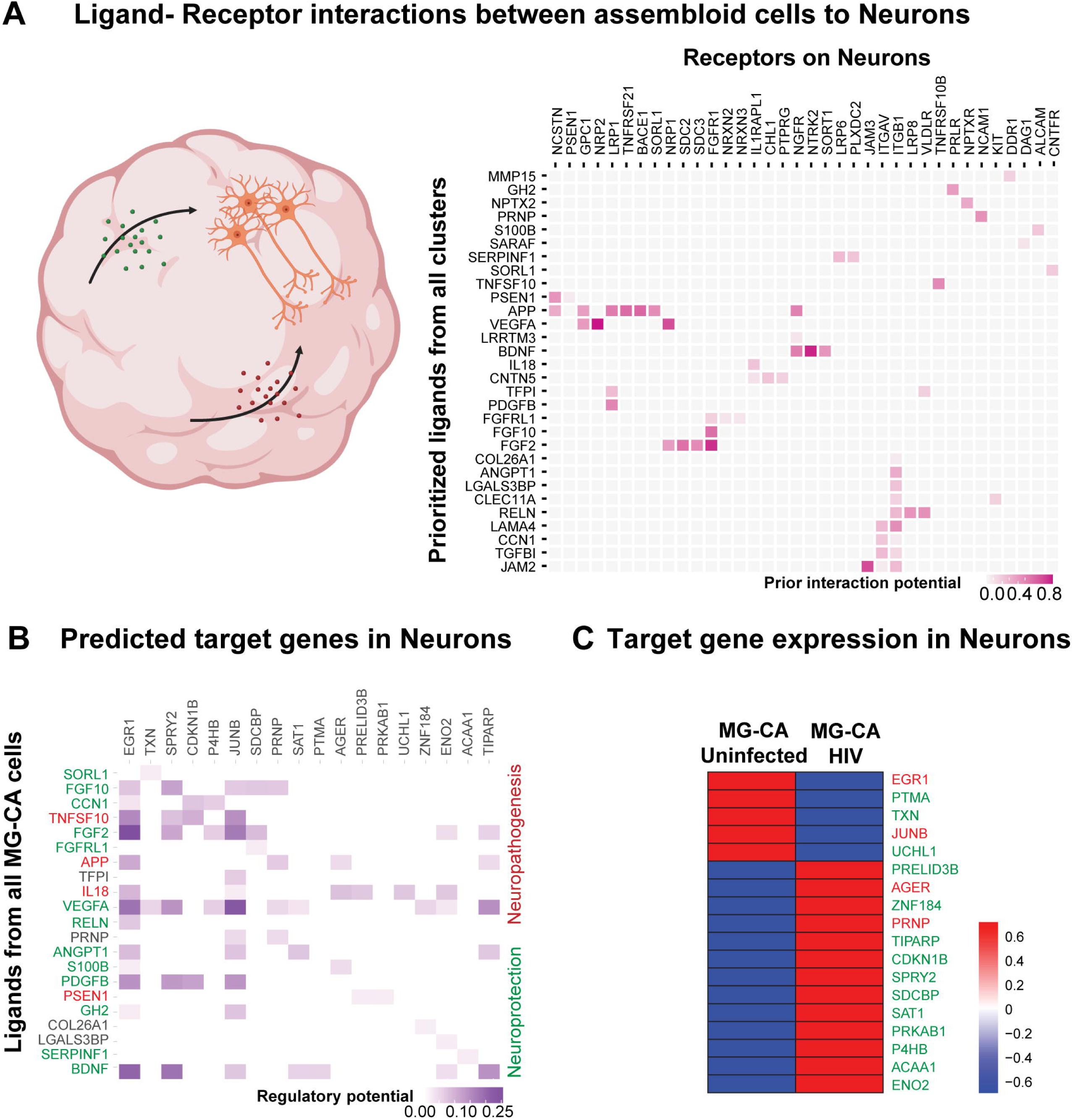
HIV infection enhances neuronal antigen presentation and stress phenotypes. **A)** Volcano plot comparing differentially expressed genes (DEGs) between neurons from MG-CAs_HIV and MG-CAs_Uninfected. The x-axis represents average log2 (fold change), while the y-axis shows-log10 (p-value). Genes with a log2 (fold change) > 0.5 and p < 0.01 are highlighted with red dots. **B)** Volcano plot showing DEGs from activated pathways identified between neurons from MG-CAs_HIV and MG-CAs_Uninfected (from **Supplementary Table 6**). The x-axis represents average log2 (fold change), while the y-axis shows-log10 (p-value)

Interestingly, we observed increased expression of genes involved in the ER stress response pathway, including those encoding ER chaperones, components of ER-associated degradation (ERAD) for aggregated proteins, and regulators of autophagy **(Fig. 8B)**. Upregulated genes included *MARCHF6* and *PARK7* (which alleviate ER stress and protect against ferroptosis [95–97]), the autophagy-related gene *ATF4* that reduces cell death during acute ER stress and shields against neuronal injury [97, 98]), *DNAJB2, CLU, UBQLN1,* and *SGTA* (which is involved in proper protein folding and clearance of misfolded proteins [97, 99]) (**Fig. 8B**).

The upregulation of gliogenesis and anti-apoptotic pathways was also observed in neurons during HIV infections. Genes within these pathways included neuroprotective genes such as PTN, DAG1, MDK, PTPRZ1, UBB (Ubiquitin B, which aids in the degradation of misfolded proteins) [100], *PRDX2* encoding peroxiredoxin 2, an antioxidant protein that protects cells from oxidative stress [101], *TRIAP1*, Heterogeneous nuclear ribonucleoprotein K (HNRNPK) that supports cell survival by inhibiting the activity of various caspases. [102] (**Fig. 8B**). Together, these gene signatures suggest cellular efforts to reduce ER stress and highlight compensatory mechanisms in neurons to alleviate ER stress and sustain homeostasis.

### Neuroinflammatory and neuroprotective signals in the HIV-infected assembloid microenvironment affect neuronal responses

To better understand the role of the assembloid environment in influencing neuronal responses during HIV infection, we mapped potential ligands from all assembloid cell types to receptors on neuronal clusters. Inflammatory ligands such as TNFSF10, APP, IL18, were among the top interactors, which are also known to be associated with neuroinflammation and degeneration. [103–105] (**Fig. 9A**). Neurotrophic protective ligand-receptor pairs were also identified, including BDNF-NTRK2, LRRTM3-NGFR, VEGFA-NRP1/2, APP-NCSTN, and FGF2/FGF10-FGFR1 **(Fig. 9A)**. Our analysis revealed that neuroinflammatory ligands such as TNFSF10 (TRAIL), IL18, and APP are involved in activating stress-responsive downstream target genes in neurons, such as EGR1 and JUNB. Conversely, neuroprotective ligands like BDNF, VEGFA, PDGFB, FGF2, FGF10, and RELN are associated with activating some homeostatic and protective genes related to redox balance, DNA repair, and cell survival, including TXN, ENO2, ACAA1, SPRY2, and CDKN1B **(Fig. 9B, left panel)**.

**Figure 9:**
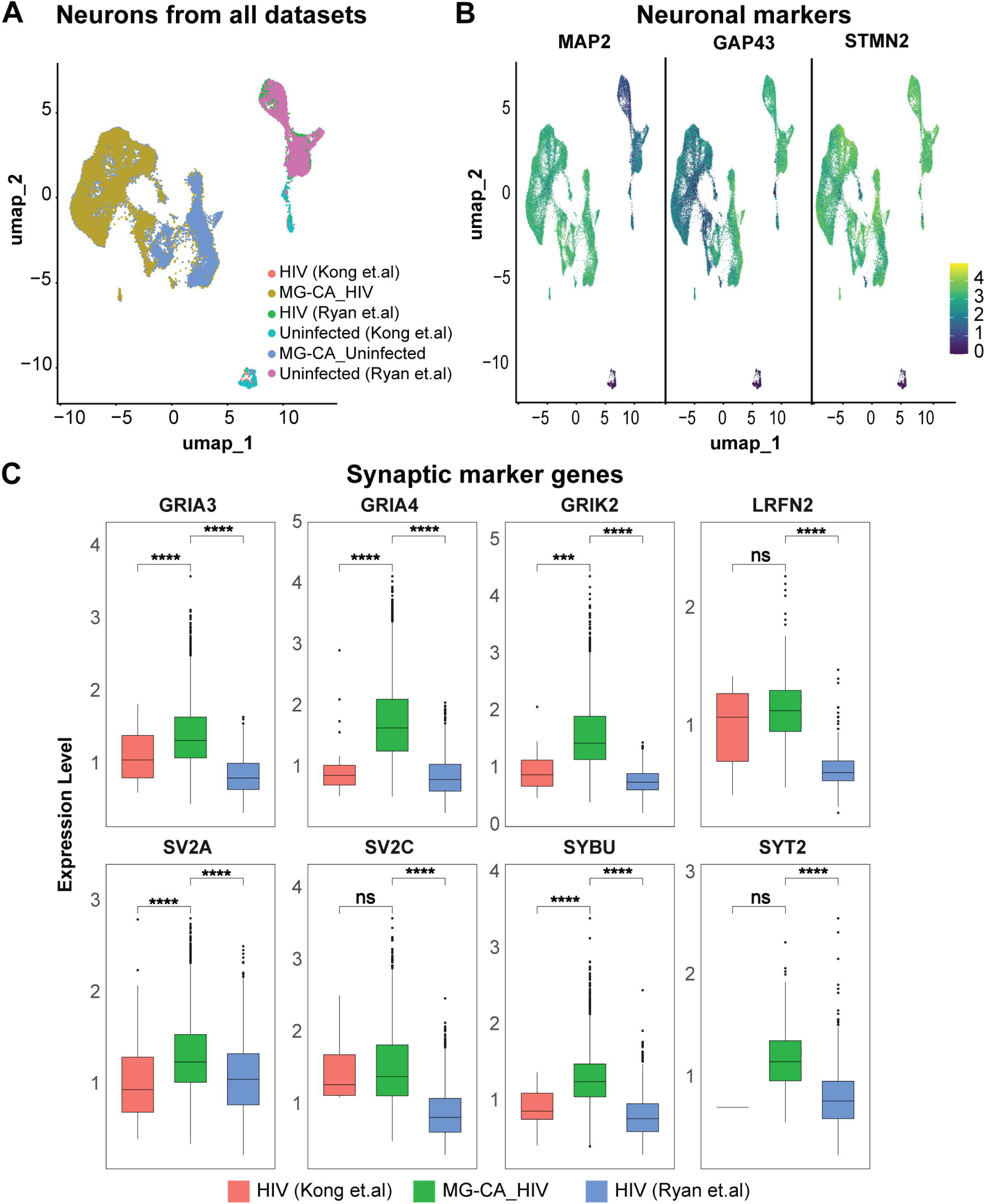
Pro-Inflammatory conditions in HIV-infected assembloids induce stress and repair responses in neurons. **A)** NicheNet interactome analysis predicting top ligand-receptor interactions between MG-CA_HIV environment cell types and neurons. The y-axis represents potential top ligands expressed by all cell types in assembloids, while the x-axis shows their cognate receptor interactions with neurons. Interaction strength is scaled in pink. **B)** NicheNet interactome analysis predicting the regulatory potential of downstream target genes in neurons through ligand-receptor interactions between MG-CAs_HIV microenvironment cells and neurons. Potential top ligands from the assembloid cells are shown on the y-axis and downstream target genes in neurons on the x-axis, with the strength of regulatory potential colored in violet. **C)** Heatmap showing the average expression of the downstream target genes in neurons from MG-CAs_HIV and MG-CAs_Uninfected. Scale shows z-scored average expression, with red indicating high expression, blue indicating low expression. Genes involved in neuroprotective, resolution of inflammation, and repair functions are highlighted in green, while genes associated with pro-inflammatory and neuroinflammatory responses are highlighted in red.

An analysis of the downstream target gene expression levels in neurons revealed the combined effects of these signaling responses. It showed a relative decrease in early genes involved in neurodegeneration, such as EGR1 and JUNB [106, 107] (highlighted in red) in MG-CA_HIV compared to MG-CA_Uninfected. Conversely, we observed a relative increase in the expression of genes that promote neuroprotection and neuronal survival under stress, such as SPRY2, ENO2, ACAA1, SAT1, SDCBP, TIPARP, and PRKAB1 (highlighted in green) in MG-CA_HIV **(Fig. 9C, right panel)**. The sources of these ligands varied by cell type. Neurons and OPC-like cells mainly expressed ligands such as VEGFA, APP, NPTX2, and RELN. In contrast, HIV RNA+ microglia showed relatively higher levels of pro-inflammatory ligands (TNFSF10, IL18, S100B) and immunoregulatory factors (GH2, SERPINF1, TGFBI) compared to bystander microglia, which exhibited increased expression of neurotrophic ligands such as BDNF and CNTN5 **(Fig. S3A-B)**. Thus, HIV infection disrupts neuronal homeostasis through microglial immune signaling, triggering stress responses, injury, and repair mechanisms in neurons. These results highlight that, in this system—where homeostatic microglia are present in the assembloids before HIV infection, along with other glia and non-glial cells—a complex interaction occurs among injury, damage, and adaptation responses during the early stages of HIV neuropathogenesis.

### Impact of the HIV-induced inflammation on other non-glial cells in the assembloids

Upon mapping the myelination and oligodendrocyte lineage pathways from the GSEA database onto neuronal populations in the assembloids, we observed that oligodendrocyte differentiation pathways were significantly enriched in the MG-CAs_HIV group compared to MG-CAs_Uninfected **(Supplementary Table 7)**. This analysis allowed us to further investigate the OPC-like population in the assembloids and track the ligands from OPC-like cells that influence all cell types in the MG-CA during HIV infection. We detected an increased potential for neurogenerative factors such as FGF2 and ephrin family genes (EFNBP), as well as ligands involved in neuronal synaptic plasticity and maintenance (NRG3, NEGR1, CNTN2, CNTN4, JAM3). These ligands showed higher interaction potential with corresponding receptors on non-glial cell types like astrocytes, neurons, and NPCs **(Fig. S4A)**. Additionally, these ligands from the OPC-like population were relatively upregulated under MG-CAs_HIV conditions **(Fig. S4B)**. These gene expression patterns align with the ability of OPC-like cells to promote axon growth, synaptic remodeling, and neuroprotective responses within the assembloid environment.

To identify the specific astrocytic subpopulations impacted by HIV in the MG-CA, we created a volcano plot comparing astrocytic clusters **(Fig. 4A)** from MG-CAs_HIV and MG-CAs_Uninfected. The plot displays average log2 (fold change) versus log10 (false discovery rate or FDR) for all genes. Genes that showed a twofold upregulation or downregulation with an FDR less than 0.01 are marked with red dots. Some of the upregulated genes included those related to the MHC class I antigen presentation family, such as CD74, B2M, HLA-B, and HLA-C, as well as genes involved in chemokine responses (e.g., CCL22) and interferon responses (e.g., IRF9, IFITM1) in astrocytes **(Fig. S5A)**.

To gain deeper insights into the cellular crosstalk affecting astrocytic reactive states in MG-CA_HIV, we performed a NicheNet analysis to examine ligand-receptor interactions between assembloid cell types and the astrocytic cluster **(Fig 4A)**, as well as to predict downstream genes activated in astrocytes from ligand-receptor signaling during HIV infection. This analysis revealed that TNF, TNFSF12, and IL-18 ligands from the HIV-infected assembloid environment activate numerous downstream NF-κB and IRF genes in astrocytes. Interestingly, neuroprotective ligands from assembloid cells—such as PDGFs, KITLG, VEGFA, and TGFB1—induce expression of ADORA2B, SPRY2, ID3, and ITGA5 genes in astrocytes, which are associated with anti-inflammatory and immunomodulatory effects. [108–110] **(Fig. S5B)**. The relative expression of these protective genes, along with inflammatory genes, was also increased in astrocytes from MG-CAs_HIV compared to those from MG-CAs_Uninfected, suggesting the presence of compensatory pathways and mechanisms that help regulate inflammation and promote cell survival **(Fig. S5C)**.

To identify the source of these ligands that affect astrocytes in the MG-CAs_HIV, we analyzed the relative expression of ligand genes across all assembloid cell types. This analysis showed that HIV-infected microglia impact astrocytes by releasing proinflammatory ligands, including IL18, LTB, and TNFSF12. Meanwhile, other cell types, such as neurons, OPC-like cells, and NPCs, expressed ligands necessary for neural development functions **(Fig. S5D)**. These findings suggest that while HIV-infected microglia trigger immune responses and MHC activation in astrocytes, neighboring non-glial cells communicate with astrocytes, influencing their inflammatory and reactive states and helping them manage inflammation-mediated effects.

### Neuronal integrity is reduced after prolonged HIV infection

To gain insights into how our neuroHIV assembloid model compares with published models, we compared neurons from our MG-CA_Uninfected and MG-CA_HIV models alongside neurons from HIV-infected choroid plexus organoids from Kong et al. [60] and the triculture model from Ryan et al. [49]. While Kong et al. documented neuronal death and dysfunction during HIV infection [60], Ryan et al. focused on microglial activation and inflammatory signatures [49]. For our comparative analysis, we used reference samples from both studies that characterized cells with ongoing viral replication, without ART suppression, for about one month after infection. In contrast, our model focused on the early infection stages (i.e., at 3 dpi) and viral dissemination.

Neuronal cells from all datasets were identified by mapping to the STAB brain reference [75], and selecting clusters with neuronal gene signatures as illustrated in **Fig. 4**. Using Seurat’s integration function followed by Harmony analysis to correct batch effects, we merged all neuronal datasets. UMAP clustering revealed clear separation along UMAP_1, indicating significant transcriptional differences between neurons from our assembloid and those from Kong et al. and Ryan et al. **(Fig. 10A)**. Expression of neuron-specific markers such as MAP2, GAP43, and STMN2 further confirmed the neuronal identity in all datasets **(Fig. 10B)**.

**Figure 10:**
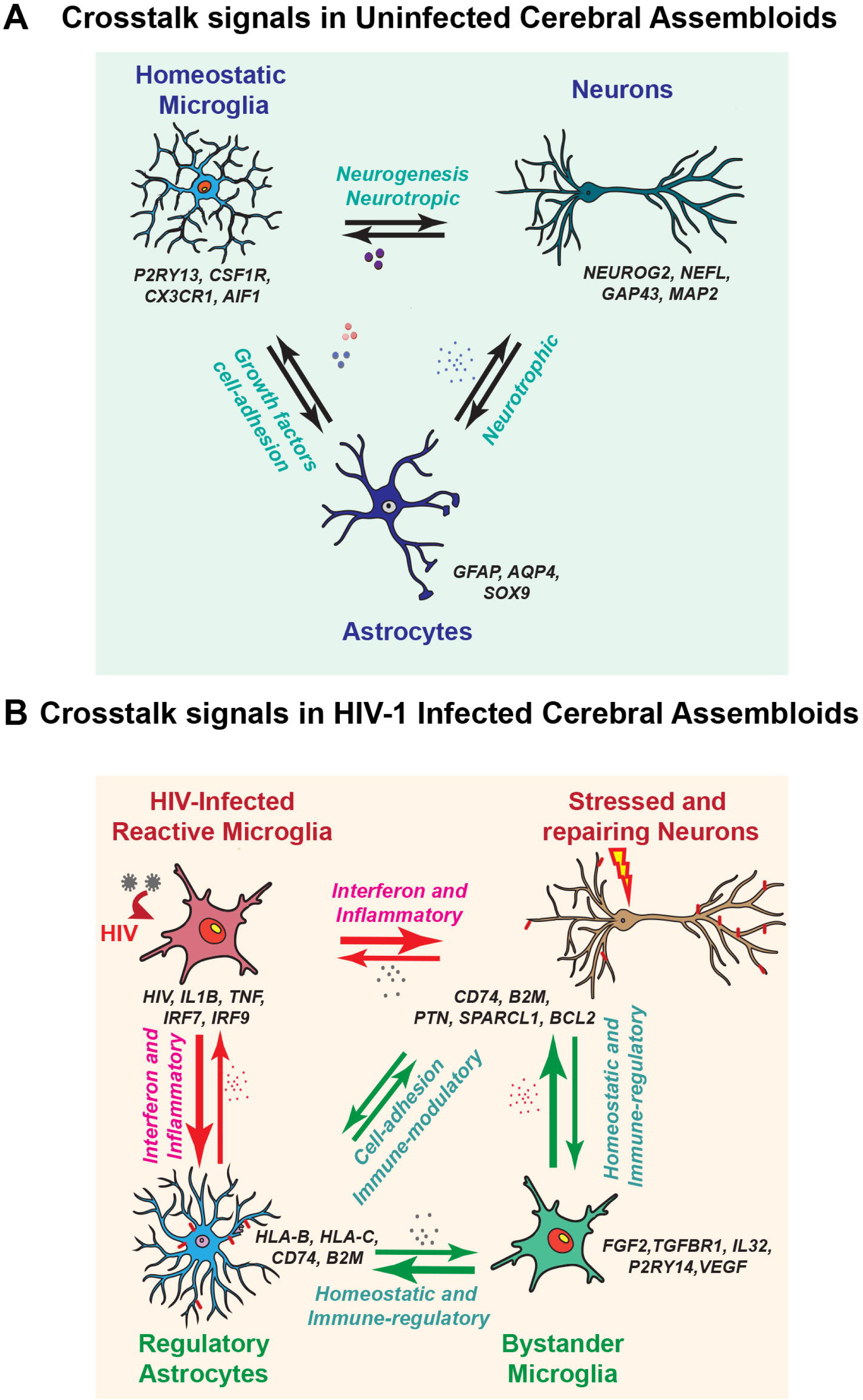
Comparison of neuronal population with reference datasets. **A)** UMAP plots showing unsupervised clustering of neuronal cells from all datasets. HIV (Kong et.al) and Uninfected (Kong et.al) represent neurons from choroid plexus organoids, HIV (Ryan et.al) and Uninfected (Ryan et.al), indicate neurons from triculture model and MG-CA_ Uninfected and MG-CA_HIV are neurons (Ex-Neurons and Inh-Neurons) from our dataset. **B)** Feature plots showing expression of neuronal markers *MAP2*, *GAP43* and *STMN2* in all populations confirming neuronal identity. **C)** Box plots show the relative expression levels of synaptic marker genes (*GRIA3, GRIA4, GRIK2, LRFN2, SV2A, SV2C, SYBU,* and *SYT2*) in neurons from all datasets; HIV (Kong et al.) (red), MG-CA_HIV (green), and HIV (Ryan et al.) (blue). Statistical significance between groups was determined using a Wilcox test with pairwise comparison; *ns* denotes non-significant differences, while asterisks indicate significance levels (p < 0.05, p < 0.01, p < 0.001, p < 0.0001).

A combined GSEA analysis across all three models revealed upregulation of inflammatory, neutrophil migration, chemotaxis, and immune response pathways in neuronal data sets from both Kong et al. [60] and Ryan et al. [49]. In contrast, neurons in our MG-CA_HIV exhibited enhanced activation of ER stress responses, as well as glial cell genesis and glial cell differentiation pathways (**Supplementary Table 8**).

We also observed that our neurons maintained elevated levels of homeostatic and neuronal identity markers such as MAP2, INA, synaptic genes (SYN1, SYN2), neuron-microglia communication markers (CD200, CX3CL1, NRXN2), mature neuronal marker RBFOX3 (NeuN), GAP43, NCAM2, and inhibitory neuron markers (GAD1, GAD2) compared to neurons from both reference datasets **(Fig. S6A)**. In contrast, neurons from the HIV-infected triculture model [49] showed increased inflammatory markers (*IL1B*, *CCL4*, *CXCL8*), while neurons from the HIV-infected choroid plexus organoids [60] exhibited increased B2M and caspase genes (CASP1), indicating heightened inflammatory stress and neuron apoptosis in the late stages of infection **(Fig. S6B)**.

Additionally, neurons in both MG-CA_Uninfected and MG-CA_HIV maintained relatively higher levels of markers indicating synaptic integrity, such as glutamate receptor subunit genes (GRIA3, GRIA4, GRIK2) and genes for synaptic vesicle proteins (SV2A, SV2C, SYT2), which are also linked to cognitive and synaptic stability. [111] **(Fig. 10C, Fig S5C)**.

To further assess neuronal responses during HIV infection, we conducted a DEG analysis comparing neurons from our MG-CA_HIV samples with neurons from Kong et al.’s HIV-infected organoids **(Supplementary Table 8)** [60]. Consistent with their findings, we observed upregulation of mitochondrial and ATP synthase genes, as well as S100 calcium-binding proteins, in their neuronal datasets [60] (**Supplementary Table 8**). In contrast, neurons from our MG-CA_HIV model maintained expression of neuronal genesis genes, including MAP2, MATR3, and MAP6. GSEA pathway analysis also showed that our neurons preserved pathways for neurite differentiation, axon development, synapse assembly, and neuronal maturation compared to neurons from Kong et al. [60] (**Supplementary Table 8**). Gene-level comparisons showed that after HIV infection in the Kong et al. model [60], the neurons showed increased expression of neuronal death markers (CASP1, PERP, PMAIP1), inflammatory cytokines and chemokines, and MHC components **(Fig. S6D)**. In contrast, our neurons from MG-CA_HIV showed higher expression of genes related to cell survival, repair, gliogenesis, neuronal generation, and ER stress regulation **(Fig. S6D)**.

These comparative analyses suggest that early HIV infection, characterized by reduced viral replication, does not cause widespread neuronal apoptosis or death. Instead, there is activation of stress management, repair, and compensatory pathways, along with signals from the microenvironment to counteract inflammation-induced damage and maintain neuronal stability **(Fig. 11A-B)**. However, as infection advances and viral replication increases, these protective mechanisms may become overwhelmed, leading to neuronal dysfunction and neurodegeneration.

**Figure 11:**
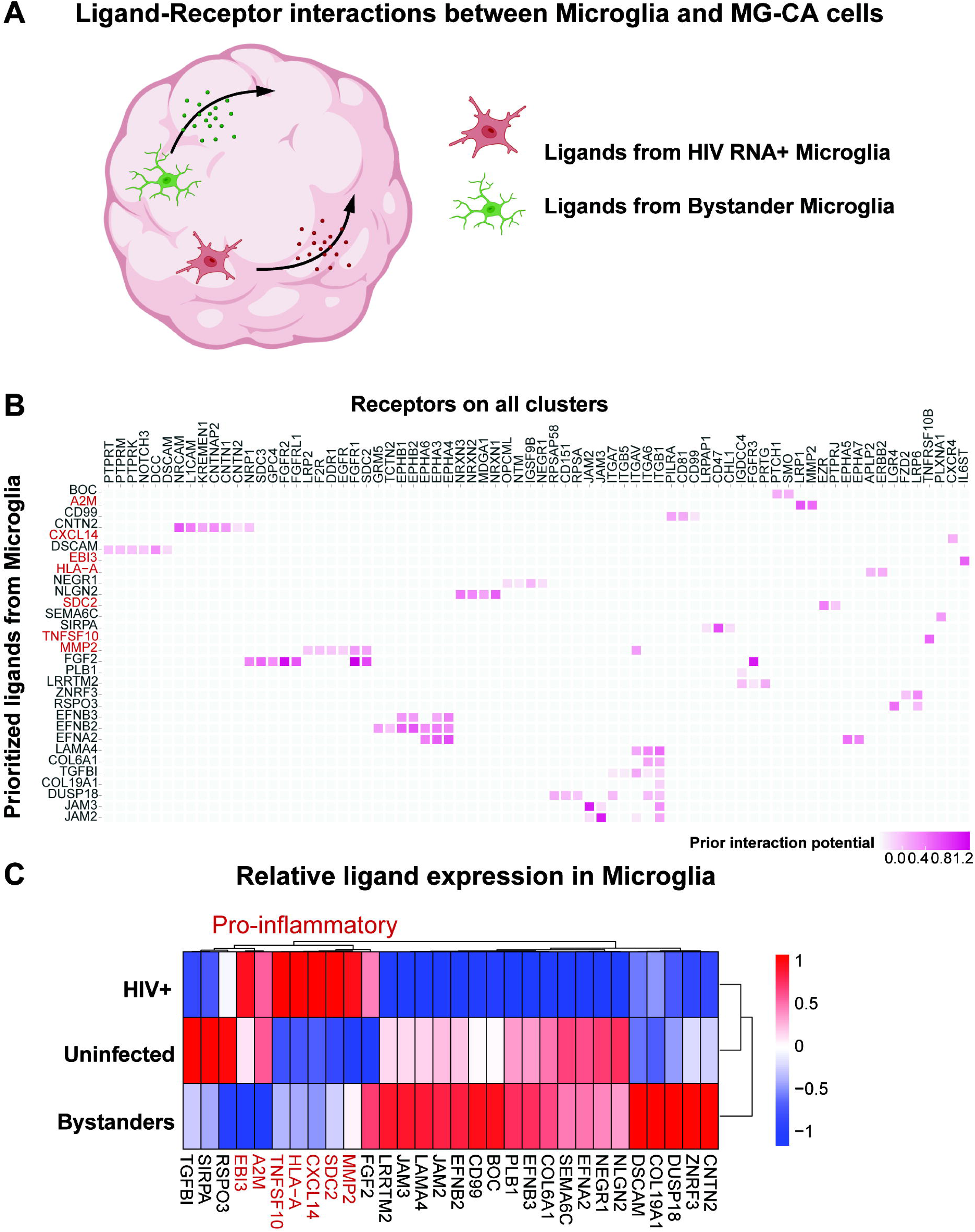
Homeostatic interactions in microglia-containing human cerebral assembloids combat early-stage effects of HIV-1 replication and inflammation. In steady-state conditions in the brain assembloids, homeostatic factors from microglia such as BDNF, NLGNs, and TGFβ1 support neuron and glial/non-glial cell differentiation, maintenance, and maturation, promoting neuroprotective phenotypes. HIV-1 primarily infects and spreads within the microglial cells in the brain assembloids. During the early acute stages of HIV infection, HIV-infected microglia exhibit a reactive phenotype with upregulated pro-inflammatory and interferon genes and reduced homeostatic genes. Reactive microglia further relay inflammatory signals to the assembloid environment, shifting neurons and other non-glial cells towards a stressed state, characterized by upregulation of MHC class I and stress response genes. Compensatory and regulatory, repair signals from bystander microglia, other glial and non-glial cells aim to counteract pro-inflammatory conditions by attempting neural repair. These events highlight the complex interplay between viral replication, inflammation, and host responses during very early stages of HIV infection, indicating mechanisms that potentially prime neurons, astrocytes, and non-glial cell types toward a neurodegenerative phenotype.

## Discussion

### Establishment and characterization of a unique assembloid model for NeuroHIV

A variety of experimental strategies have been used to incorporate microglia into cerebral organoids to model CNS HIV infections [49, 56–61]. Despite notable progress, challenges persist in replicating the homeostatic interactions of the brain microenvironment [60, 61, 76], sustaining long-term microglial viability, high inter-organoid variability, and mimicking the developmental origins of microglia [49, 57, 58]. Adapting the co-development method described by Xu et al. [62], we created a microglia-containing cerebral assembloid system by combining iPSC-derived tdTomato+ microglial precursors with tdTomato-negative NPCs. Key modifications included adding an endogenous tdT tracker to visualize microglial differentiation, partially pre-differentiating hematopoietic stem cells toward the microglial lineage, and expanding the NPC before assembloid formation. This allowed for faster microglial differentiation alongside the development of neuronal and other non-glial cells. By day 15, tdT+/IBA-1+ microglia appeared at physiologically relevant levels, exhibiting diverse phenotypes related to homeostasis, neurogenesis, phagocytosis, chemo-sensing, and cytokine responses, which are characteristic of early developmental and prenatal microglia [25, 27]. We observed that microglia influence neuronal development through ligands such as NLGNs, NEO1, BDNF, and TGF-β1, which support synaptic function and maturation. These findings establish our MG-CA system as a reliable platform for studying homeostatic regulation and the effects of HIV infections on neuroimmune interactions.

### Effect of HIV infection on assembloid microenvironment

Another key advance in our study is that, unlike previous models that pre-infect microglia before incorporating them into organoids [49, 57, 58], or focus on late-stage HIV effects and characterize global transcriptomic changes using bulk RNA-sequencing [60, 61], we introduced HIV particles directly into the assembloid culture media and examined the early effects of HIV on individual cell types within the assembloids. This approach mimics natural infection and captures the initial responses and interactions between HIV and the brain microenvironment, similar to CNS viral seeding during peripheral infection.

In our model, microglia are the primary targets of HIV, capable of harboring the replicating virus. Although the assembloids contained an astrocyte-like cell population, CO_HIV showed no detectable proviruses or HIV RNA. HIV replication persisted up to 5 dpi, after which proviral levels began to decline, likely due to virus-induced toxicity and the death of infected cells. Notably, we observed an increase in spliced HIV mRNA from 3 to 4 dpi, which gradually decreased by five dpi as infected cells either died or probably entered latency. Supervised clustering identified two distinct microglial populations within HIV-infected assembloids (MG-CA_HIV): productively infected microglia (HIV RNA+) and bystander microglia (HIV RNA = 0). Phenotypic differences showed that bystander microglia had higher expression of homeostatic genes and pathways compared to HIV RNA+ microglia. Conversely, HIV RNA+ microglia exhibited a reactive inflammatory phenotype and significantly modulated the microenvironment by releasing interferon-responsive and pro-inflammatory ligands. Microglial immune signaling disrupted neuronal homeostasis, triggering stress responses, HLA gene expression, injury, and repair pathways in neurons and other non-glial cells. These findings underscore the impact of HIV infection on microglia, as well as glial and non-glial cell types within our assembloids.

### Inter-cellular interactions to resolve inflammation-mediated effects

By modeling early pathogenesis, we identified inflammatory immune activation and stress response genes in neurons and astrocytes that occur without overt neurodegeneration. Our cell-cell interaction analysis showed that, besides inflammatory signals, important ligands from the MG-CA_HIV microenvironment supported neuronal survival and repair.

Synaptodendritic injury, a well-documented feature of HAND, arises from damage to pre-and postsynaptic structures without causing significant neuronal death, and this process may be reversible through neuronal plasticity [112, 113]. Neuronal plasticity allows neurons to recover function after brain injury and is characterized by increased dendritic branching, axogenesis, formation of new synaptic connections, and some neurogenesis [3, 94, 114]. Key mediators from the brain microenvironment and coordinated crosstalk signals among microglia, neurons, astroglia, and OPCs can influence synaptodendritic plasticity and repair after injury. [3, 115–122].

A recent multi-omic study by the Spudich group [123], identified dysregulated microglial interactions in PWH, especially pro-inflammatory signaling. Analogously to our findings, analysis of interactions among microglia, astrocytes, and neurons showed upregulation of synaptic organization and neuronal migration factors (NRXN1, LAMA4, NLGN1, BDNF, VEGFA, WNT5A, FGF2), as well as immune-related interactions (B2M, HLA-E, ITGAV, LRP1, NRP1/2) in PWH. These shared signatures suggest synaptic changes linked to HIV, possibly as an adaptive response to neuroinflammatory damage.

## Conclusions

The regulatory mechanisms aimed at restoring homeostasis and balancing injury and repair during HIV infection may determine the repeated cycles of viral reactivation and suppression in the brain [42]. Our previous research showed that healthy neurons and anti-inflammatory agents can suppress HIV transcription in microglia. [42–47]. Alternatively, the ongoing interaction of low-level viral replication, neuronal stress, and inflammatory signaling, along with repair efforts, may delay neurotoxicity and synaptic dysfunction but can become overwhelmed as HIV continues to spread. Comparing this with late-stage infection models that lack viral suppression helps us understand how these repair and compensatory mechanisms may fail, leading to significant neurodegeneration. Disruptions in the balance between injury and repair signals likely also increase the severity of cognitive symptoms in HAND [3]. Our analysis of the early stages of infection using the assembloid systems helps us rigorously define the balance between injury and adaptation responses and provide valuable insights for developing therapies that enhance neuroprotective pathways, target microglial activation, and reduce HIV reservoirs in the brain. In agreement with Ellis et al. [3], our results strongly suggest that supplementing ART during HIV infections with neuroprotective agents may help to facilitate CNS repair and recovery.

## Materials and Methods

### Reagents and Materials

**Supplementary Table 9** shows a list of all antibodies, reagents, deposited data, and software used in this study.

### Maintenance and generation of tdTomato-tagged iPSC lines

The iPSC line used in this study was generated from a neurotypical male individual (“Clay” from Marchetto et.al [124]). The reprogramming and validation of this cell line were previously described [124]. In a subsequent study, the genome of this cell line was edited via CRISPR/Cas9 and TALEN systems to incorporate components of the COR-LT lineage tracing system (“control COR-LT” from Bury et al. [64]). Among these components was a “STOP-flox” nuclear-localized tdTomato reporter that was integrated into the AAVS1 safe harbor site [125, 126]. To generate a tdTomato-positive cell line for the current study, the “control COR-LT” cell line was transiently transfected with cre recombinase, which removed the STOP sequence between the tdTomato reporter and CAG promoter in a subset of transfected cells. This enabled permanent expression of tdTomato in these cells. To establish a pure tdTomato+ population, transfected cells were plated at clonal density, isolated after 3-5 days in culture, and further expanded. Fluorescent imaging was used to confirm tdTomato expression in all cells from clones used to generate organoids in this study. Human iPSC line CS00iCTR-n2, described in **Fig. 1**, was obtained from Cedars-Sinai Medical Center’s David and Janet Polak Foundation Stem Cell Core Laboratory (CS vial ID# 1017819, source: male, fibroblast). All iPSC lines were validated for pluripotency markers OCT3/4, TRA-1-60, SOX2 and confirmed normal karyotype.

iPSC lines were maintained in complete TeSR-E8 medium (Stem Cell Technologies) until confluency and detached as small aggregates using ReLeSR (Stem Cell Technologies). The aggregates were centrifuged at 1000rpm for 1 minute. The pellet was resuspended in stem cell medium plus ROCK inhibitor Y-27632 (RI;10 mM) and plated on 6-well tissue culture plates coated with gelatin (Thermo Fisher). The cells were incubated at 37°C in a 5% CO2 incubator. Geltrex coating was performed by mixing aliquots of geltrex, thawed on ice, with ice-cold DMEM medium at a 1:100 dilution, followed by incubating 1 mL per well for 1 hour at 37°C in 5% CO2.

### Generation of tdTomato-tagged hematopoietic stem cells (iHSC) and microglia (iMGs) from iPSCs

To generate tdTomato-tagged iHSCs, clay tdTomato iPSCs were differentiated into hematopoietic stem cells using the STEMdiff hematopoietic kit (Stem Cell Technologies). Briefly, iPSCs were dissociated with ReleSR into small aggregates measuring between 100 μm and 150 μm and plated at approximately 80-100 aggregates per well in a 6-well plate with E8 media containing RI. The total number of colonies plated was carefully selected for each iPSC line after testing different colony dilutions per well and assessing yields of CD34+ cells. After confirming the optimal size and number of colonies the next day (10-20 colonies/cm²), E8 media was replaced with fresh media A at days-16 and-14 of iHSC differentiation. To initiate the second phase of cell differentiation, media A was replaced with media B at day-13. From day-11 until day-4, cells received fresh media B every other day. Starting on day-9, round, floating iHSCs became visible. On day-4, at the end of the 14-day iHSC differentiation process, floating iHSCs were harvested through gentle pipetting and characterized for hematopoietic stem cell markers CD43, CD34, and CD45 by flow cytometry **(Supplementary Table 1)**.

To induce iMG differentiation from iHSC, we followed the original protocol by the Blurton-Jones group [127]. The day (-4) iHSC were transferred to microglia (iMG) media (as described in Abud et.al [65]) containing 100 ng/mL IL-34, 50ng/mL TGFβ1 and 25ng/mL M-CSF. iHSCs at day 1 (5 days post transfer to iMG media) were used to generate microglia-containing organoids.

### Generation of neural progenitor cells (NPCs) from iPSCs

The protocol for generating neuronal progenitor cells through neural induction was outlined by Gibco (Life Technologies). In brief, control untransfected COR-LT iPSC lines without tdTomato expression were seeded at a density of 2.5 x 10^5^ cells per well in a 6-well plate using Gibco Neural Induction Medium, which includes 490 mL of Neurobasal medium with 10 mL of neural induction supplement, with daily medium changes. By day (-22) during neural induction, the cells reached near-maximal confluence, and any non-neural differentiated cells were removed. On day (-23), neural stem cells (NSCs) at passage P0 were harvested and expanded in neural expansion medium, containing 49 mL of Neurobasal medium with 2 mL of neural induction supplement and advanced DMEM/F12, up to passage P4 NSCs.

### Generation of cerebral organoids and assembloids containing microglia

To generate microglia-containing assembloids, NSCs P4 cells were dissociated using TrypLE (Thermo Fisher) and combined with hematopoietic (microglial) precursors. A mixture of 70% NSCs P4 cells and 30% hematopoietic (microglial) precursors was plated at a density of 1 x 10^4 cells per well in V-shaped 96-well plates, using 50% NPC medium and 50% microglia medium. To generate cerebral organoids without microglia, 100% NSCs P4 cells were plated in the same way. The NPC medium contained 237 mL neurobasal medium, 237 mL DMEM/F12, 5 mL N2 supplement, 10 mL B27 supplement minus vitamin A, 1% non-essential amino acids (NEAA, Gibco), 1% GlutaMAX, 1% Penicillin-streptomycin, and 100 ng/mL FGF2. The microglia medium contained 250 mL DMEM/F12 with Glutamax, 2x insulin-Transferrin-Selenium, B27 serum-free, N2 supplement, NEAA, and 5 µg/mL human insulin. Finally, all organoids and assembloids were maintained for up to 15 days in NPC and microglia medium supplemented with 100 ng/mL IL-34, 50 ng/mL TGFβ1, and 25 ng/mL MCSF.

Starting from day 15, NPC II medium was used to maintain all organoids and assembloids. The NPC II medium consisted of 250 mL neurobasal medium, 250 mL DMEM/F12, 5 mL N2 supplement, 20 ng/mL BDNF, 20 ng/mL GDNF, 200 nM ascorbic acid, 0.4 mM cAMP, 100 ng/mL IL-34, 25 ng/mL MCSF, and 50 ng/mL TGFβ1. After 25 days, 100 ng/mL CD200 and 100 ng/mL CX3CL1 were added to the NPC II medium.

### 2D immunocytochemistry on organoid/assembloid sections

On the indicated days after starting the organoids and assembloids (Day 15 and Day 25), they were fixed with 4% (wt/vol) formaldehyde in Tris-Buffered Saline (TBS) for 1 hour, washed with TBS, and then submerged in TBS with 30% sucrose overnight. The organoids and assembloids were mounted in 50% OCT embedding compound and 50% 30% sucrose, frozen in dry ice, and stored at-80°C. To prepare sections for staining, 20µm thick slices were cut with a cryostat and thaw-mounted onto negatively charged histological slides. The slices were stored at-20°C for up to six months. Before staining with antibodies, the slides were thawed at room temperature for 10 minutes and then submerged in TBS for 5 minutes. Antigen retrieval was performed using an antigen retrieval buffer containing 10 mM sodium citrate and 0.05% Triton X-100 (pH 7.4) twice, each for 10 minutes. To block non-specific binding, 5% donkey serum in TBS was applied for 1 hour at room temperature. Primary antibodies were applied overnight at 4°C, and secondary antibodies were applied for 45 minutes at room temperature.

### 3D immunocytochemistry of whole organoids/assembloids

To prepare the organoids and assembloids for staining, we followed the same fixation process as the 2D immunocytochemistry protocol. After fixation, the organoids and assembloids were kept in TBS with 30% sucrose overnight and then transferred to Fructose-Glycerol (FG) medium containing 60% glycerol and 2.5M fructose for clearing [128]. For staining, the entire organoid and assembloid were incubated with primary antibody for 24-48 hours at 4°C on a shaker, followed by incubation with the secondary antibody for 24 hours at 4°C on a shaker. DAPI was added for 30 minutes at room temperature on a shaker. Finally, the whole organoids and assembloids were loaded onto negatively charged histological slides, covered with mounting medium, gently covered with cover slips, and dried overnight at 4°C before imaging. Confocal microscopy was performed using a Leica TCS SP8 gated STED 3X microscope with a 10× objective, a numerical aperture of 0.4, and z-sectioned at 2 µm intervals.

### DNA-free HIV-1 preparation and pNL4-3-based DNA contamination assay

To produce DNA-free stocks of replication-competent, macrophage-tropic HIV-1, we transfected 293T cells in 6-well plates with 4□µg per well of the pNL(AD8) plasmid [129, 130]. To improve viral titers and eliminate pNL(AD8) DNA, we collected culture supernatants 2 days after transfecting 293T cells, which were treated with DNase I, and used them to infect CEMx174 5.25 cells (provided by Nathaniel Landau). These cells are human lymphoid cell lines engineered to stably express human CCR5 and a GFP reporter under the control of the HIV-1 5’ LTR promoter, so only infected cells express GFP. The infected cells were then cultured in RPMI 1640 medium with 10% FBS until more than 40% of the cells were GFP-positive. These cells were washed with PBS, resuspended in microglia media at a density of 1 million viable cells per mL, and cultured for an additional 48 hours. The resulting culture supernatant was harvested, clarified by centrifugation (1500 x g for 5 minutes), filtered through a 0.45 µm filter, and stored in aliquots at-80°C. The specific infectivity of this stock was confirmed by infecting 5.25 cells and detecting GFP-positive cells via flow cytometry within 2 days. Viral fitness was verified by an increase in the percentage of GFP-positive cells at 7 days post-infection. The level of pNL(AD8) plasmid contamination in 1 µL of virus was measured using digital PCR with primers and probe targeting the pNL4-3 DNA sequence outside the HIV-1 proviral genome, including pNL4-3 backbone reverse (pNL4-3_9719-9739), HIV-1 LTR forward (HXB2_500-522), and HIV-1 LTR TaqMan VIC-MGB probe (HXB2_533-560). Viral titers of each inoculum were determined using the Lenti-X™ GoStix Plus kit (Takara).

### HIV Infection of organoids and assembloids

To induce HIV infection, we added replication-competent NLAD8 (∼130 ng p24) directly into cell culture media on day 15 in 96-well plates. To improve viral penetration into organoids and assembloids, we incubated the plates on an 80 rpm shaker at 37°C. We continued monitoring the cultures without media changes for up to 3 days post-infection (dpi) and performed half-media changes every other day after 3 dpi until 6 dpi.

### RNA and DNA extraction

DNA and RNA isolation followed the Qiagen protocol (AllPrep DNA/RNA Mini Kit). Organoids and assembloids were resuspended in Buffer RLT Plus (supplied with the AllPrep DNA/RNA Mini Kit) and vortexed to lyse the cells. The lysate was directly passed into a QIAshredder (Qiagen) spin column placed in a 2 mL collection tube and centrifuged for 2 minutes at full speed. The flowthrough was collected and passed through the QIAshredder again to ensure complete homogenization of the organoids and assembloids. Lysates from organoids and assembloids were stored in RLT buffer at-80°C until processing. RNA and DNA extraction were performed according to manufacturer instructions (Qiagen), and concentrations were measured using Qubit™ dsDNA HS (High Sensitivity) and Qubit™ RNA High Sensitivity (HS) Assay Kits as directed by the manufacturer.

### HIV proviral assay

Proviral loads were measured in Microfluidic Array Plates (MAP16) using the QuantStudio Absolute Q Digital PCR System (Applied Biosystems). 300-700 ng of cellular DNA per well was mixed with Absolute Q Digital PCR Master Mix and the following primer/probe sets: 900 nM gag forward primer (HXB2_1282-1305), 900 nM gag reverse primer (HXB2_1347-1367), 900 nM env forward primer (HXB2_6510-6534), 900 nM env reverse primer (HXB2_6563-6586), 250 nM TaqMan® gag FAM-MGBNFQ probe 3’ (HXB2_1307-1326), and 250 nM TaqMan® env VIC-MGBNFQ probe (HXB2_6537-6560). Digital PCR conditions included 96°C for 10 minutes, followed by 40 cycles at 96°C for 5 seconds and 60°C for 15 seconds. Fluorescence was measured using the Absolute Q Digital PCR Instrument and analyzed with the QuantStudio Absolute Q Digital PCR System Software. Data from multiple wells are pooled during analysis to improve confidence. The number of human cells in each PCR mixture was calculated based on the cellular DNA concentration and the mass of the human diploid genome (6.5 × 10^-12^ g DNA per cell).

### Spliced HIV-1 mRNA Assay

To measure spliced HIV RNA, 10 ng of total cellular RNA was combined with Absolute Q RT-PCR Mix, 900 nM of the 5’LTR forward primer (HXB2_612-635), 900 nM of the tat/rev reverse primer (HXB2_5977-5996), and 250 nM of the TaqMan® FAM MGB HIV psi probe (HXB2_685-708). The PCR mixes were loaded into MAP16 digital PCR plates following the manufacturer’s instructions. RT-PCR was performed at 55□°C for 10 minutes, then at 96□°C for 10 minutes, followed by 40 cycles at 96□°C for 5 seconds and 60□°C for 10 seconds. Using the QuantStudio System software, threshold signals in the FAM channel were determined based on samples with uninfected cell RNA, and the copies of target RNA per microliter detected were converted to copies per nanogram based on input RNA concentration.

### Dissociation of organoids/assembloids for scRNA-seq

Organoids and assembloids were dissociated into single cells using the Worthington Papain Dissociation System kit, following a previously described protocol (https://protocolexchange.researchsquare.com/article/pex-258/v1, 31) with the following modifications. About 20 pooled organoids and assembloids were used per condition. The organoids and assembloids were removed from culture medium and washed once with PBS. After the wash, PBS was removed, and papain/DNase solution was added to initiate dissociation, as described in (https://protocolexchange.researchsquare.com/article/pex-258/v1, 31). Once dissociation was complete, cells were diluted in PBS with 0.04% BSA (Millipore Sigma), counted for viable cells using trypan blue, and kept on ice until processed for scRNA-seq cell capture.

### Single-cell RNA sequencing (scRNA-seq)

scRNA-seq was performed using the BD Rhapsody HT Express platform and the Whole Transcriptome Analysis (WTA) Amplification and Library Preparation Kit (BD catalog # 666620) following the manufacturer’s instructions. Briefly, approximately 60,000 cells were suspended and loaded onto the BD Rhapsody cartridge. Single cells settled in microwells with barcoded beads. With an estimated cell capture efficiency of about 70%, we expected to capture roughly 40,000 single cells. After lysis, polyadenylated mRNA was captured by the oligo(dT) tails of barcoded oligonucleotides covalently attached to the beads. Following reverse transcription and treatment with Exonuclease I, cDNA libraries were prepared and sequenced on the NovaSeq X 10B next-generation sequencing platform (Illumina, Inc) at Medgenome, Inc., using 0.5 lanes per library with a sequencing depth of over 40,000 reads per cell.

### Bioinformatic analysis

Raw fastq files were subjected to quality control using FastQC software (https://www.bioinformatics.babraham. msigdbr ac.uk/projects/fastqc/). After uploading to the Seven Bridges server (https://igor.sbgenomics.com), raw reads in fastq files were mapped using BD Rhapsody Sequence Analysis Pipeline. We used human GRCh38 primary assembly genome, GeneCode v.42 primary assembly annotation, and HIV-1 full-length genome (HXB2, GenBank: K03455.1) [93] as references, along with the reference tdTomato indices. Resulting gene expression count tables were used to create single-cell experiment objects using the Seurat package for R [131]. After filtering out low-quality cells with a suboptimal number of UMI (*DropletUtils* package for R [132]), Our yield was over 10,000 high-quality single-cell profiles per sample, with more than 8,000 UMI per cell and at least 1000 genes per cell. Gene expression counts were log-normalized, scaled, and variable features and principal components were identified. Significant principal components were determined using ElbowPlot and Jackstraw analyses. This was followed by finding neighbors, clusters, and UMAP dimensionality reduction of the first twenty significant principal components. Cell cycle stages of the single cells were estimated using the CellCycleScoring function in Seurat. Cell types were predicted by computational mapping to a brain reference dataset [75, 133] and further classified into subsets using RNA assays [134]. Gene expression data in the objects was denoised to remove ambient RNA contamination, including background HIV reads.

Denoising of the datasets was performed using a Bayesian algorithm with the DecontX package for R (doi:10.18129/B9.bioc.decontX). Denoised reads were used to perform supervised clustering shown in Fig. 5 to identify the HIV RNA+ population and prevent false positive reads caused by alignment errors. The Seurat objects from different samples were merged and integrated, minimizing batch effects with the Harmony package for R. [135] The batch effects were corrected after integrating two independent replicates or reference datasets, followed by standard object processing that included normalization, scaling, identifying variable features, PCA, finding neighbors, clustering, and UMAP reduction. UMAP images were generated using the DimPlot function in Seurat. The expression of individual genes was visualized with the FeaturePlot function. Violin plots showing differential gene expression were created using the VlnPlot function in Seurat. Differentially expressed (DE) gene markers were identified with the FindAllMarkers function in Seurat, which used the Wilcoxon Rank Sum test. The analysis was set to a Log2 fold change > 1 and min.pct = 0.1. The function searched for both positive and negative gene markers and calculated unadjusted as well as Bonferroni-adjusted p-values.

### Gene Set Enrichment Analysis (GSEA)

Gene set enrichment analysis was performed using the clusterProfiler and fgsea packages in R [136–138]. Differential expression was calculated using the FindMarkers function in Seurat with the Wilcoxon rank-sum test, applying a minimum expression threshold of 5-10% (min.pct = 0.05-0.1). The resulting gene list was ranked by average log2 fold change to create a pre-ranked gene list for GSEA. Gene sets related to Gene Ontology Biological Processes (GO:BP) were obtained from the Molecular Signatures Database [139] (MSigDB, category C5, subcategory BP) via the msigdbr package. GSEA was performed using the compareCluster function with the GSEA method, Benjamini-Hochberg correction for multiple testing (pAdjustMethod = “BH”), and gene set size filtering (minGSSize = 10, maxGSSize = 500). Results were visualized as dot plots using the dotplot function in enrichplot, showing the top enriched pathways by adjusted p-value and enrichment score.The same DE genes and reference pathway datasets, as well as the *EnhancedVolcano* package for R were used to generate volcano plots showing log_2_ fold change and p-values for the DE gene hits overlapping with the pathway gene sets (https://github.com/kevinblighe/EnhancedVolcano).

### NicheNet Analysis

Cell-cell communication analysis was conducted using the NicheNet algorithm, based on a curated database of ligand-receptor interactions, ligand-target regulatory potential matrix, and weighted signaling and regulatory networks obtained from the Zenodo repository (https://zenodo.org/record/7074291) [92]. This computational method identifies and ranks ligand-receptor pairs based on their Pearson correlation coefficient derived from the transcriptomic signatures of the ‘sender’ and ‘receiver’ populations, predicting the ability of ligands to induce a set of target genes in ‘receiver’ cells. We implemented a sender-focused approach to identify ligands from MG-CA or CO cell types that potentially influence gene expression changes in receiver cells across different conditions. Genes expressed in sender and receiver cell types were defined using a minimum expression threshold of 5%. Differential gene expression in receiver cells was calculated between the condition of interest (e.g., MG-CA_HIV) and the reference condition (e.g., MG-CA_Uninfected) using the FindMarkers function in Seurat. The set of interest consisted of differentially expressed genes (adjusted p < 0.05, |log_₂_FC| ≥ 0.25). NicheNet’s ligand activity predictions were ranked based on the area under the precision-recall curve (AUPR) and filtered to include only ligands expressed in sender populations. Top-ranked ligands were used to infer target gene regulatory interactions and receptor-ligand interaction networks. Heatmaps visualized ligand activity scores, ligand-target connections, and ligand-receptor potential. Dot plots displayed ligand expression across conditions and clusters. Controls for analyses shown in **Fig. 4** included organoids without microglia (CO_Uninfected), while MG-CA_Uninfected served as controls in **Figs. 6, 7, 9, and S3-S6**.

### Rigor, reproducibility, data analysis, and Statistics

Statistical analyses were performed using Origin. Specific statistical tests used and sample sizes (n) are indicated in the figure legends. Both technical and biological replicates were included, using two independent donor iPSC lines to ensure the reliability of the readouts. In cases of normal distributions, Student’s unpaired t-tests were used to compare pairs of groups. Otherwise, the Wilcoxon rank sum test was employed for datasets with skewed distributions. The relationships between specific markers (continuous variables) were assessed using Spearman’s correlation. Nichenet ranking of ligand-receptor pairs was performed based on the Pearson correlation coefficient.

## Declarations

### Ethics approval and consent to participate

Not applicable

### Consent for publication

Not applicable.

## Availability of data and materials

### Materials availability

All materials generated in this study will be made available from the lead contact without restriction.

### Data and code availability

Single-cell RNA-seq data generated in this study are deposited at Gene Expression Omnibus (GEO) under GSE300562 and will be made publicly available as of the date of publication. Publicly available scRNA-seq datasets from Kong. et al were obtained from GEO under accession number GSE262349. Triculture scRNA-seq datasets from Ryan et.al were obtained from the GEO accession numbers GSM4272584 and GSM4272587.

### Funding

This work was supported by grants from the NIH, NIMH (R01MH134316), NIDA (R01 DA060489 and R01 DA049481) to J.K. and A.W-B, the Rustbelt Center for AIDS Research (Case/UHC-Pitt CFAR) P30 AI036219, and NIDA CWRU Center for Excellence on the Impact of Substance Use on HIV P30 DA054557.

### Author contributions

Conceptualization, S.S., Y.C., L.A.D.B., A.W.-B. and J.K.; methodology, S.S., Y.C., L.A.D.B., K.S.L., J.E; B.L., J.H. formal analysis, S.S., Y.C., K.S.L.,; investigation, S.S., Y.C., K.S.L., and J.E.; writing – original draft, S.S., and J.K.; writing – review & editing, S.S., Y.C., L.A.D.B., K.S.L., J.E., F.Y., Y.G.M., B.L., A.K., J.K. and A.W.-B.; visualization, S.S., Y.C., and J.K.; supervision, J.K. and A.W.-B.; funding acquisition, J.K. and A.W.-B.

## Acknowledgments

We thank the Cedars-Sinai Medical Center’s David and Janet Polak Foundation Stem Cell Core Laboratory for providing the CS00iCTR-n2 iPSC line. The following reagent was obtained through BEI Resources, NIAID, NIH: Human Immunodeficiency Virus Type 1 (HIV-1) NL4-3 AD8 Infectious Molecular Clone, pNL(AD8), HRP-11346.

## Abbreviations

PWH: People With HIV
NCI: HIV-Associated Neurocognitive Impairment
ART: Antiretroviral Therapy
CNS: Central Nervous System
BBB: Blood-Brain Barrier
CO: Cerebral Organoids
ESCs: Embryonic Stem Cells
iPSCs: Induced Pluripotent Stem Cells
3D: Three-Dimensional
CA: Cerebral Assembloids
2D: Two-Dimensional
NPCs: Neural Progenitor Cells
scRNA-seq: Single-Cell RNA-Sequencing
MG-CA: Cerebral Assembloids Containing Microglial Cells
tdT: tdTomato
BDNF: Brain-Derived Neurotrophic Factor
GDNF: Glial Cell Line-Derived Neurotrophic Factor
cAMP: Cyclic-AMP
iHSC: Hematopoietic Stem Cells
STAB: Spatio-Temporal Cell Atlas of the Human Brain
iMGs: In Vitro Differentiated Microglia
dpi: days post-infection
PIPL: Potentially Intact Proviral Load Assay
MG-CA_HIV: HIV-1 NL-AD8 Infected MG-CAs
CO_HIV: HIV-Infected CO
DEGs: Differentially Expressed Genes
Ex-Neurons: Excitatory Neurons
Inh-Neurons: Inhibitory Neurons
OPC-like: Oligodendrocyte Progenitor-Like
GSEA: Gene Set Enrichment Analysis
HAND: HIV-Associated Neurocognitive Disorders
ER: Endoplasmic Reticulum
ERAD: ER-Associated Degradation

## Supplemental information

**Additional file 1: Supplementary Figures S1-S7**

**Additional file 2: Supplementary Table 1.** Characterization of iHSC markers at Day (-4) and Day 1 by flow cytometry and Immunohistochemistry (IHC) of microglia IBA-1 expression at Day 15 and Day 25 of organoid and assembloid cultures, **related to Figure 1**

**Additional file 3: Supplementary Table 2.** List of all top genes in tdT+ subclusters (clusters 0 to 7), **related to Figure 2**

**Additional file 4: Supplementary Table 3.** Quantification of HIV proviral DNA in sheet HIV_Proviral_copies and spliced HIV mRNA in sheet HIV_spliced_RNA, **related to Figure 3**

**Additional file 5: Supplementary Table 4.** Top DEGs between cell clusters from MG-CA and COs at Day 18, List of Top GSEA pathways and genes comparing neurons from MG-CA_Uninfected and CO_Uninfected, **related to Figure 4**

**Additional file 6: Supplementary Table 5.** Top GSEA pathways and genes comparing microglia from MG-CA_HIV and MG-CA_Uninfected, Top GSEA pathways and genes comparing HIV RNA+ microglia and Bystander microglia from MG-CA_HIV, **related to Figure 5**

**Additional file 7: Supplementary Table 6.** Top GSEA pathways and genes comparing neurons from MG-CA_HIV and MG-CA_Uninfected, Top GSEA enriched pathways and genes comparing neurons from MG-CA_HIV and MG-CA_Uninfected, **related to Figure 8**

**Additional file 8: Supplementary Table 7.** List of the myelination and oligodendrocyte lineage pathways from the GSEA database mapped to neurons from MG-CA_HIV and MG-CA_Uninfected, **related to Supplementary Figure 5**

**Additional file 9: Supplementary Table 8.** Top GSEA pathways and genes comparing neurons from our MG-CA (MG-CA_HIV vs MG-CA_Uninfected), Kong et.al (HIV vs Uninfected) organoids and Ryan. et.al (HIV vs Uninfected) Triculture model, Top GSEA pathways and genes comparing neurons from our MG-CA_HIV with HIV (Kong et.al) organoids, **related to Figure 10 and Supplementary Figure 7**

**Additional file 10: Supplementary Table 9.** List of Antibodies, Reagents, Deposited data and software used in this study

