## Supplementary figures and images for "Transcriptional Mapping of Neuro-Immune Interactions during Homeostasis and HIV infection using Microglia-containing Human Cerebral Assembloids"

### Supplementary Figure 1

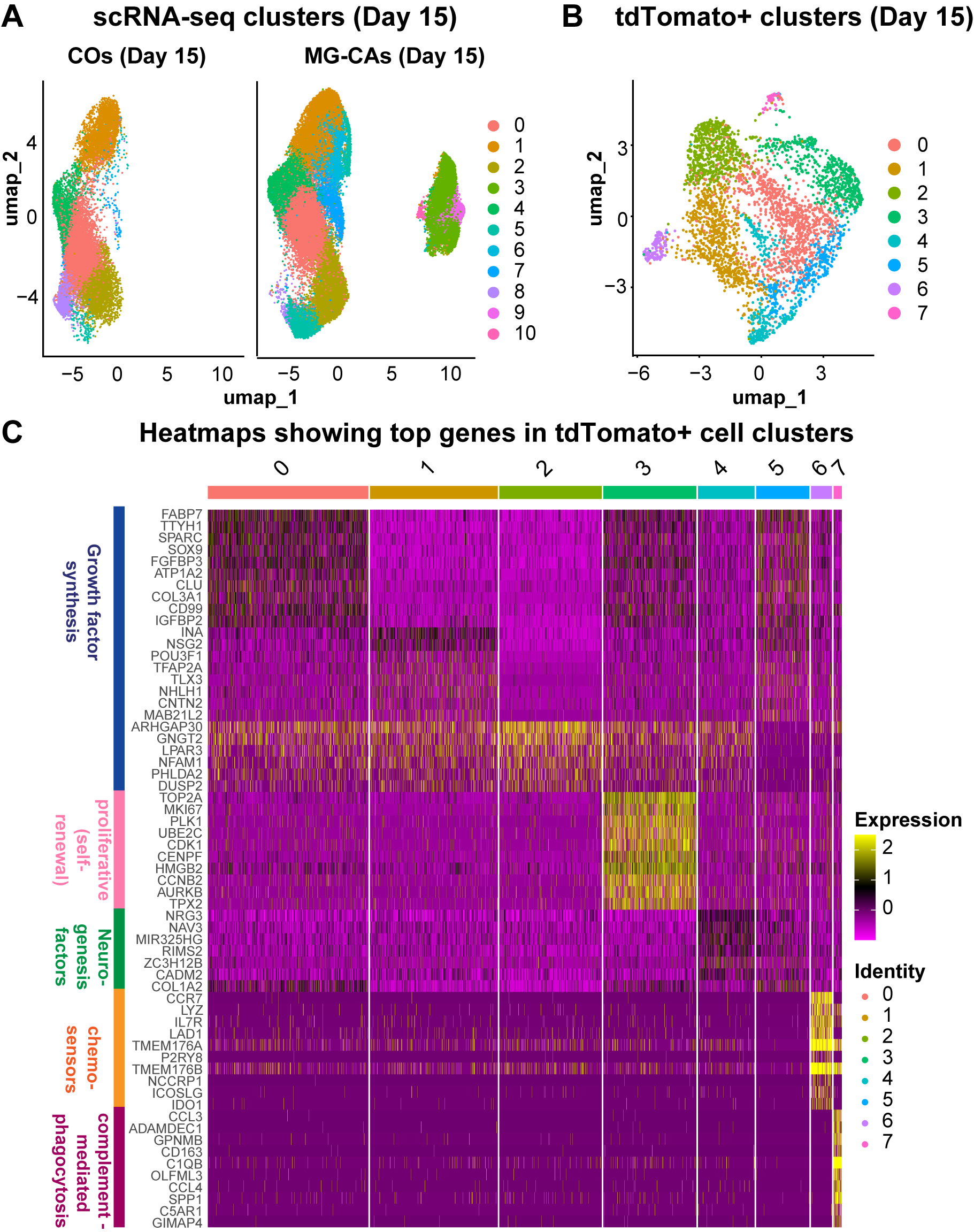

### Supplementary Figure 2

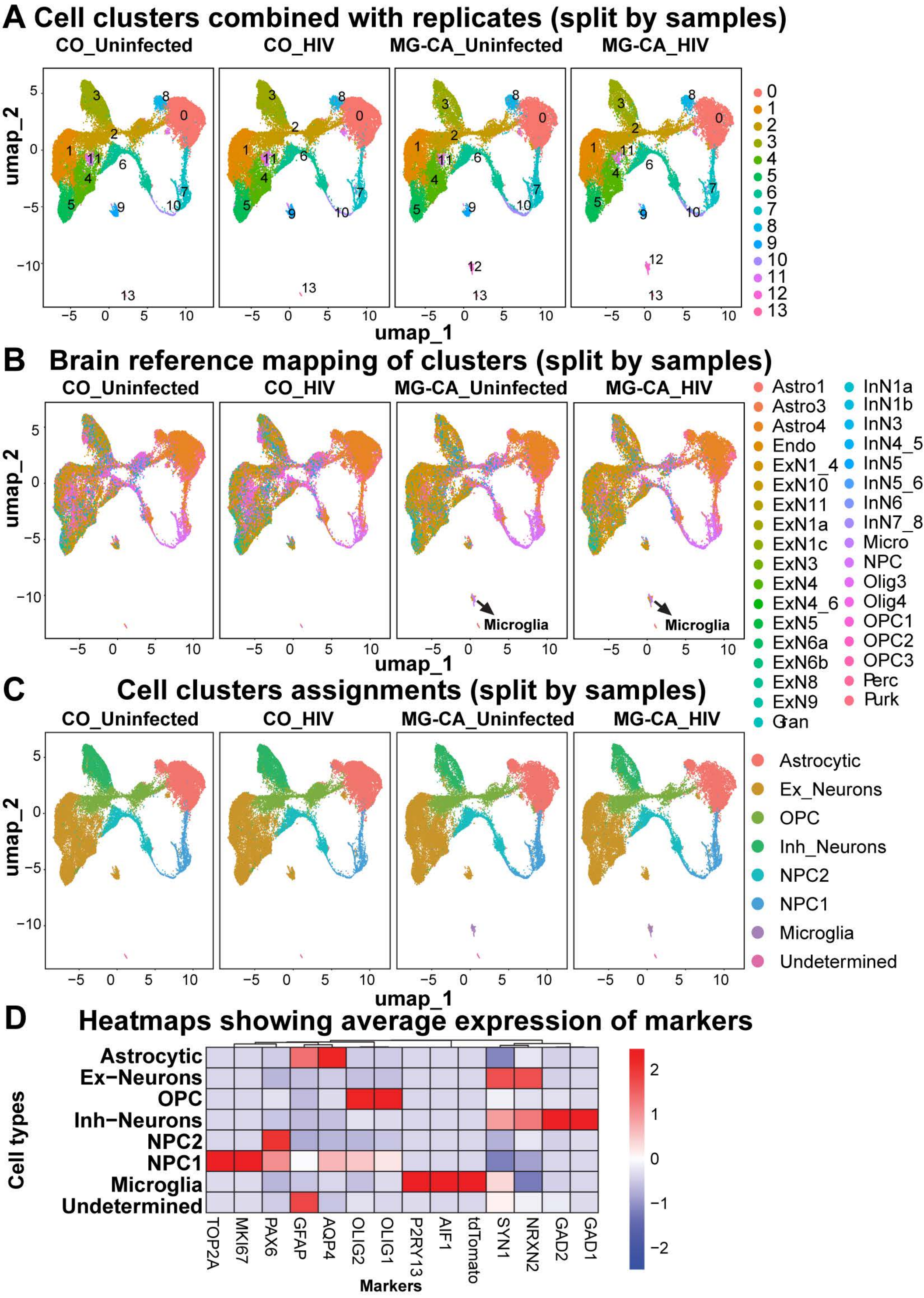

### Supplementary Figure 3

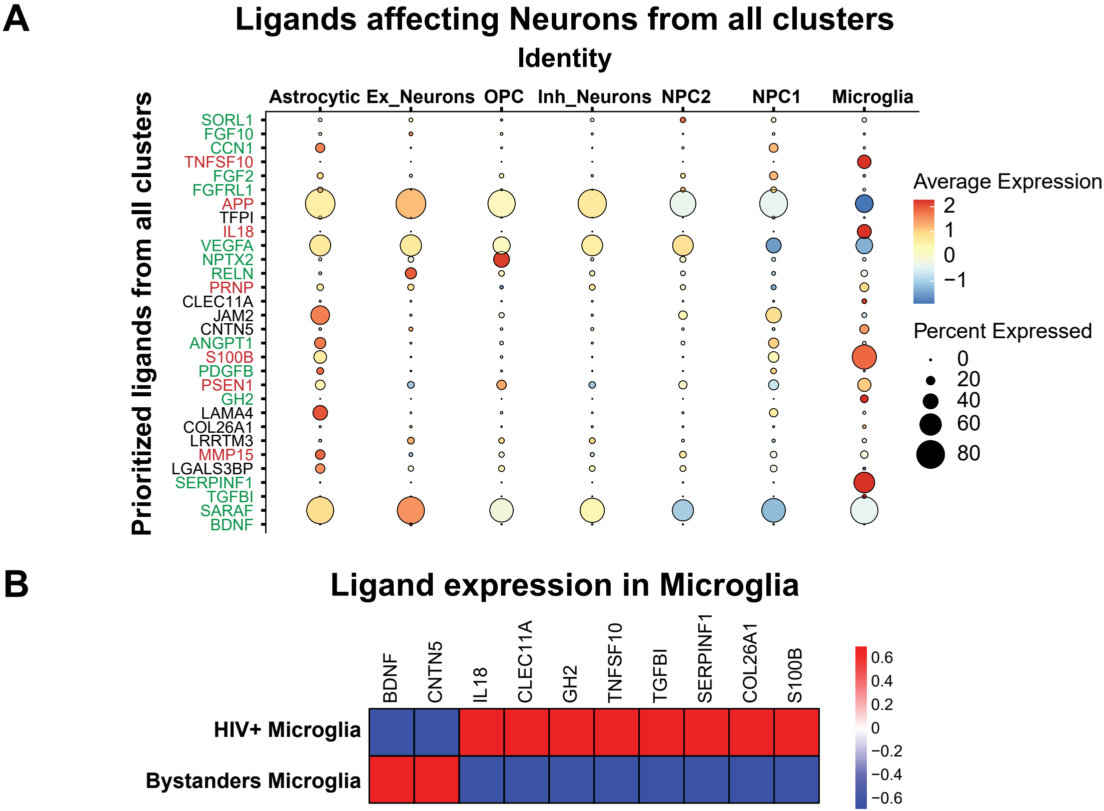

### Supplementary Figure 4

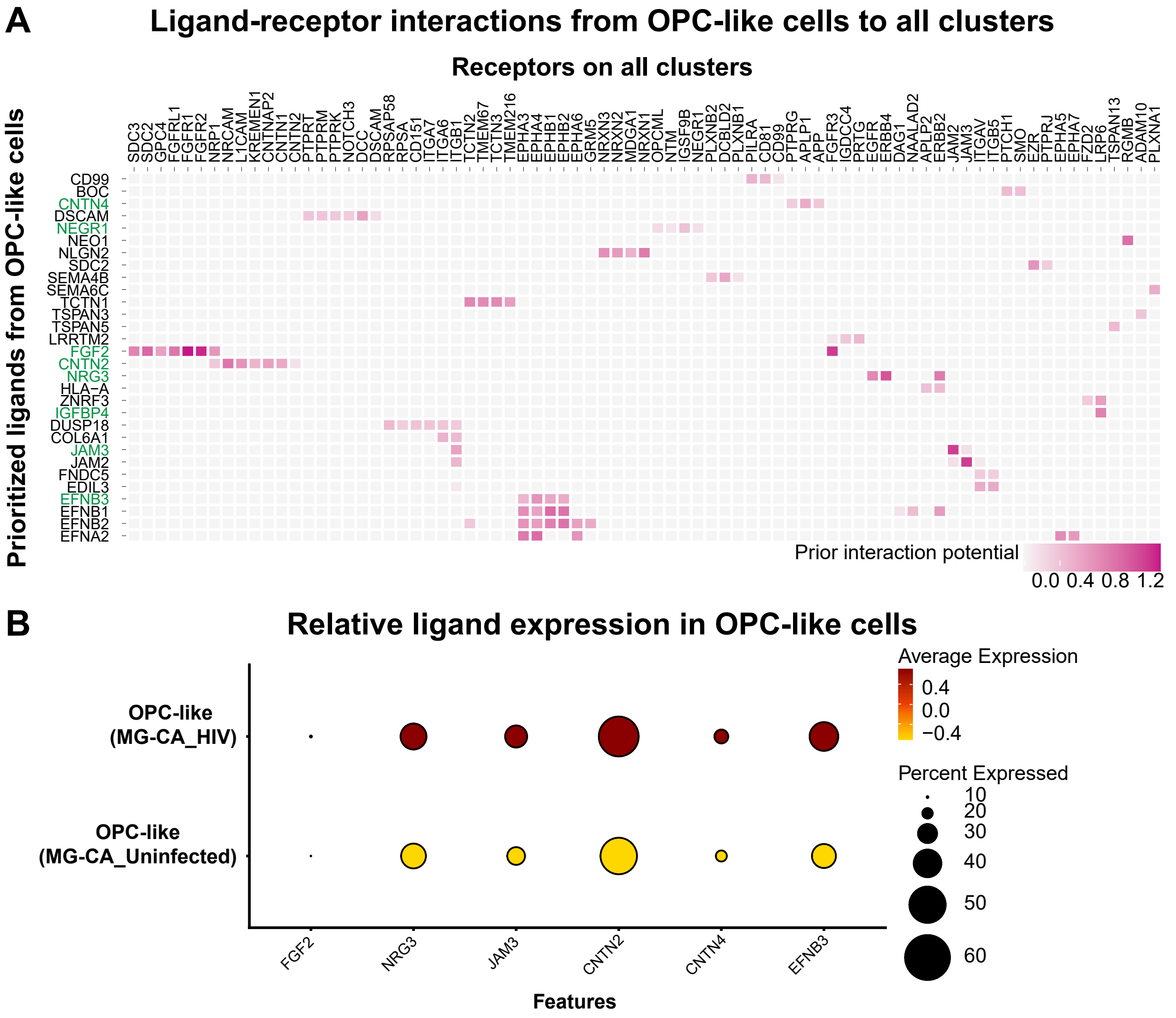

### Supplementary Figure 5

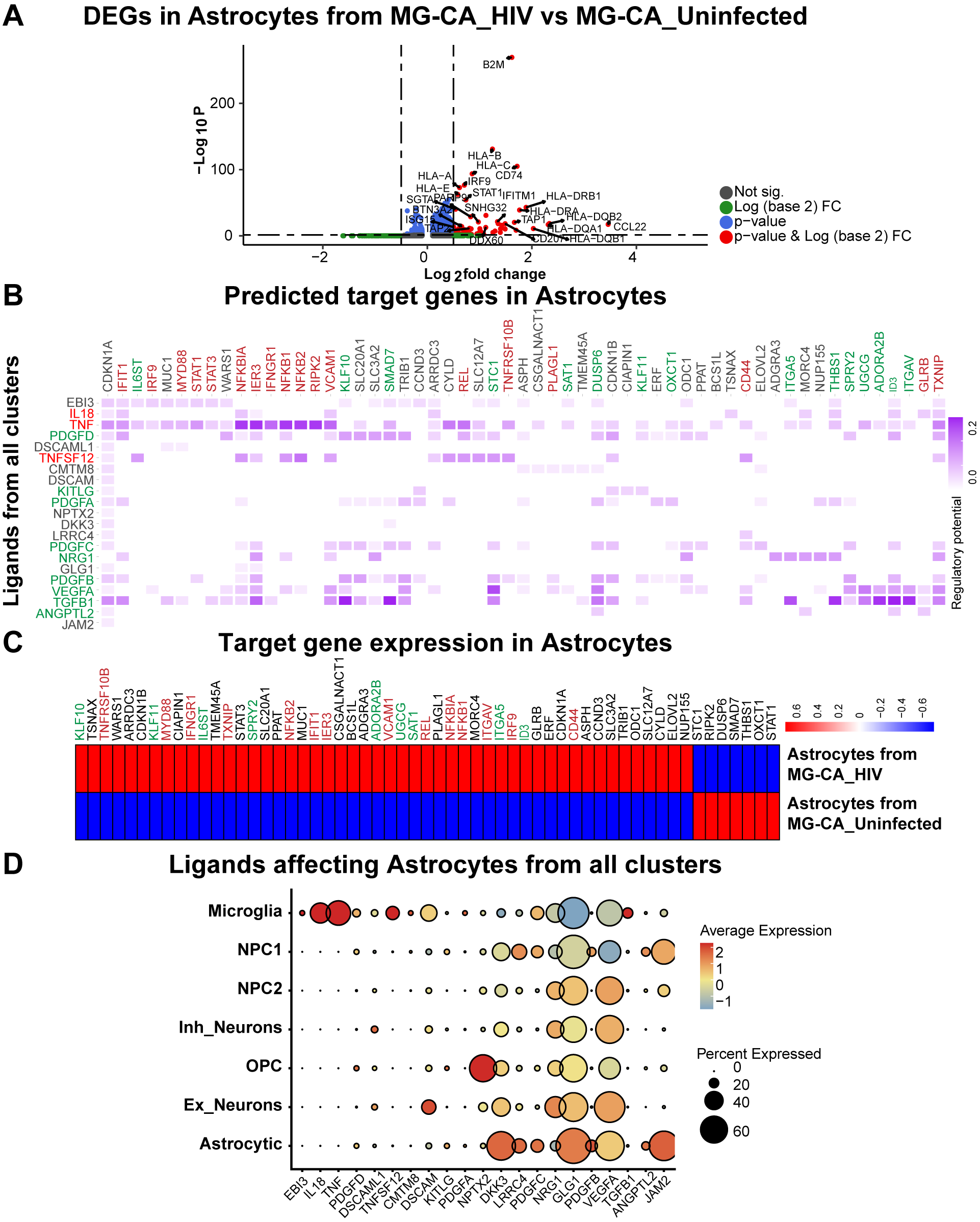

### Supplementary Figure 6

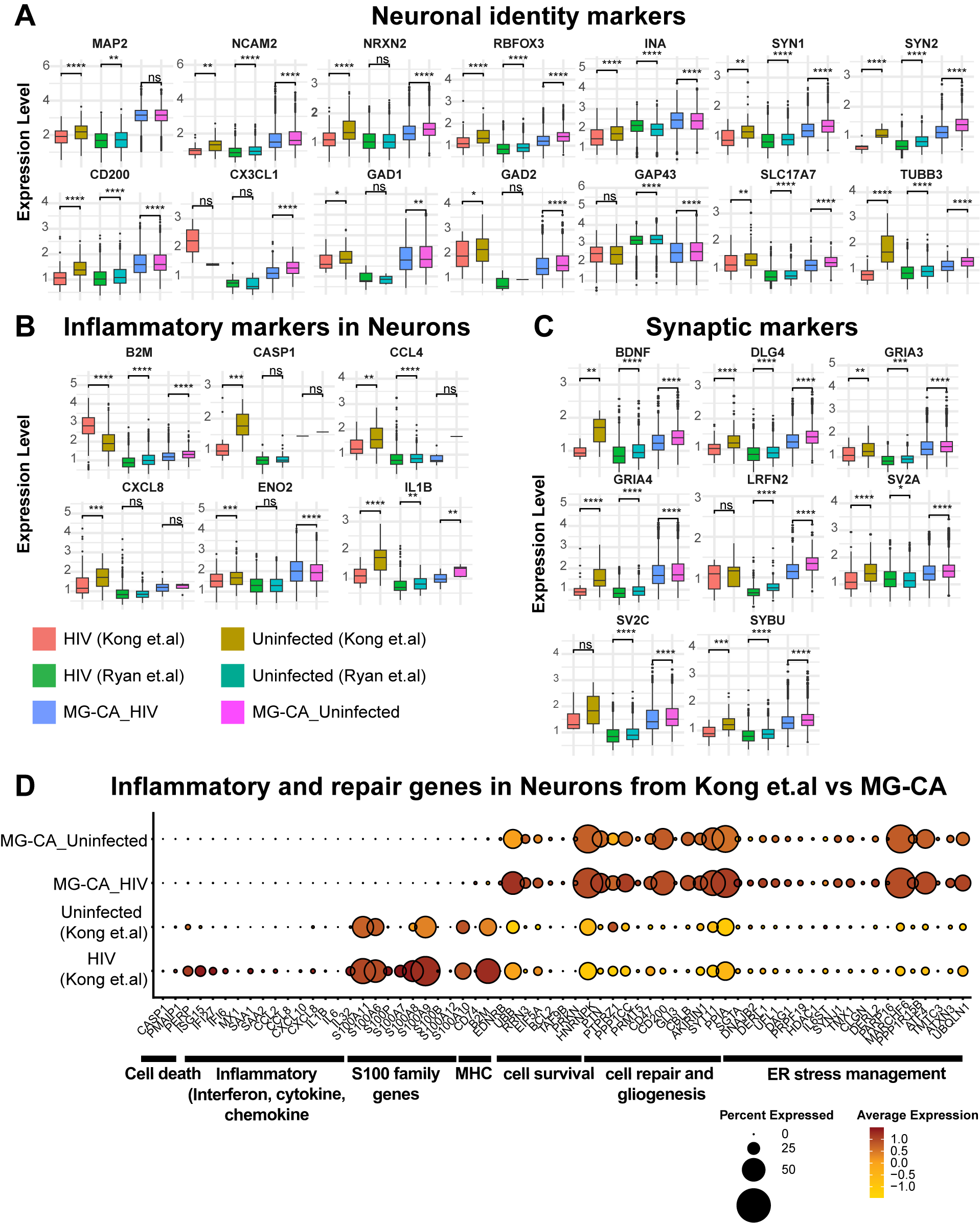
